# Universal length fluctuations of actin structures found in cells

**DOI:** 10.1101/2023.07.27.550898

**Authors:** Aldric Rosario, Shane G. McInally, Predrag R. Jelenkovic, Bruce L. Goode, Jane Kondev

**Affiliations:** Department of Physics, Brandeis University, Waltham, United States; Department of Biology, Brandeis University, Waltham, United States; Department of Electrical Engineering, Columbia University, New York, United States

## Abstract

Actin is a key cytoskeletal protein that forms filaments that bundle into linear structures *in vivo*, which are involved in motility, signaling, and cell division. Despite the rapid turnover of individual actin monomers, these structures are often maintained at a specific length, which is important for their function. Length control is commonly attributed to length-dependent assembly or disassembly of the structure, whereby a stable length is achieved when the two opposing processes are balanced. Here we show that regardless of the nature of the length-dependent feedback, such “balance point” models predict a Gaussian distribution of lengths with a variance that is proportional to the steady state length. Contrary to this prediction, a reexamination of experimental measurements on the lengths of stereocilia, microvilli, actin cables, and filopodia reveals that the variance scales with the square of the steady state length. We propose a model in which the individual filaments in bundles undergo independent assembly dynamics, and the length of the bundle is set by the length of the longest filament. This model predicts a non-Gaussian distribution of bundle lengths with a variance that scales with the square of the steady state length. Our theory underscores the importance of crosslinking filaments into networks for size control of cytoskeleton structures.

## INTRODUCTION

Actin is a major component of the cytoskeleton and is one of the most abundant proteins present in eukaryotic cells. Cells utilize actin to build a wide range of structures by polymerizing monomers into helical filaments^1^. The filaments are often then arranged into higher order, crosslinked structures that have a significant impact on cellular shape, division, motility, and signaling^2^. An important class of such structures are parallel actin bundles, which are composed of parallel filaments packed together axially by actin-bundling proteins. Filopodia at the leading edge of motile cells, stereocilia in inner and outer hair cell bundles, microvilli on the apical surfaces of epithelial cells, and actin cables in budding and fission yeast cells are a few examples of parallel actin bundles *in vivo*^3–5^.

Actin structures *in vivo* turnover rapidly, meaning that the actin monomers and their associated proteins that make up the structure are continuously being added and removed from the structure on time scales that are short when compared to the lifetime of the structure. Yet these structures often maintain a well-defined length that is critical for their function. For example, misregulation of the lengths of stereocilia in hair cells can lead to deafness^6,7^, microvilli length defects result in microvillus inclusion disease (MVID)^8^, filopodial length defects cause defects in axon guidance^9^, while actin cable defects in yeast lead to inefficient transport of endocytic vesicles which in turn produces growth deficiencies^10^.

Balance point models have been proposed as a general mechanism by which cells control the length of their cytoskeletal filaments^11,12^. The multi-step process of assembly, bundling, maintenance and disassembly of cytoskeletal filaments involves multiple proteins working in synergy with each other. Balance point models abstract this myriad of molecular processes by introducing two rate parameters, namely the rate of assembly or polymerization and the rate of disassembly or depolymerization. In the most general case, the assembly and disassembly rates are assumed to be functions of the length of the filament. This functional dependence encodes the feedback on length necessary to reach a stable, steady state length, reached when the rates of assembly and disassembly are matched, or balanced^12^.

The balance point model also describes length fluctuations in steady state. These fluctuations originate from the stochastic processes of adding and removing monomers from the filamentous structures, which transiently increase or decrease the filament length. The key question we address with this work is how measuring length fluctuation of filamentous structures in cells informs us about the molecular mechanisms of their length control.

Studying fluctuations has in the past yielded insights into a variety of biological mechanisms at play in cells^13^. A classic example is the fluctuations in the number of bacterial colonies that arise after exposure to bacteriophage, which was utilized by Luria and Delbruck to discern the mechanism by which resistance to infection arises^14^. In the context of gene expression, measurements of fluctuations reveal different mechanisms of transcriptional regulation in cells as the different mechanisms show different noise signatures^15,16^. In the study of cytoskeletal structures, measurements of length fluctuations of flagella in the unicellular algae *Chlamydomonas reinhardtii* were used recently to discriminate among the different feedback mechanisms that lead to length control^17^.

In the present study, we show that balance point models of length control predict a Gaussian distribution of length fluctuations in steady state, with a variance that scales linearly with the steady state length. This property is universal to all balance point models and does not depend on the specific functional form of the feedback. By analyzing published results on the length variation of parallel actin bundles such as stereocilia^18^, microvilli^19^, filopodia^20^, and actin cables^21^, we show that all of these structures exhibit a universal scaling of the variance with the square of the mean length, in contradiction to the predictions from balance point models.

These observations lead us to propose a model of filament bundles where we account for their crosslinked multi-filament nature. The individual filaments within the bundle undergo independent dynamics and the length of the bundle is set by the longest filament. In this case, the length distribution of the bundle derived from extreme value statistics^22,23^, leads to a peaked non-Gaussian distribution, even when filaments within the bundle are unregulated and exponentially distributed. We show that the variance of the bundle length scales quadratically with the mean length, consistent with experimental observations. Our theory emphasizes the importance of crosslinking filaments in order to produce linear structures of a specific length. Furthermore, our calculations reveal how bundle length control can arise from the architecture of the bundle without a specific feedback mechanism on the individual filaments within the bundle.

## RESULTS

### Balance point models of filament length control predict a Gaussian distribution of lengths with a variance that scales linearly with the length

Cells contain a wide range of linear structures (Figure 1A), each of which is constructed from multiple actin filaments crosslinked together (Figure 1B, left). To mathematically describe the assembly dynamics of such linear structures, we consider a long bundle of crosslinked actin filaments as an idealized polymer that grows by addition of subunits and shrinks by removal of subunits (of length *a*) (Figure 1B, right).

**Figure 1.**
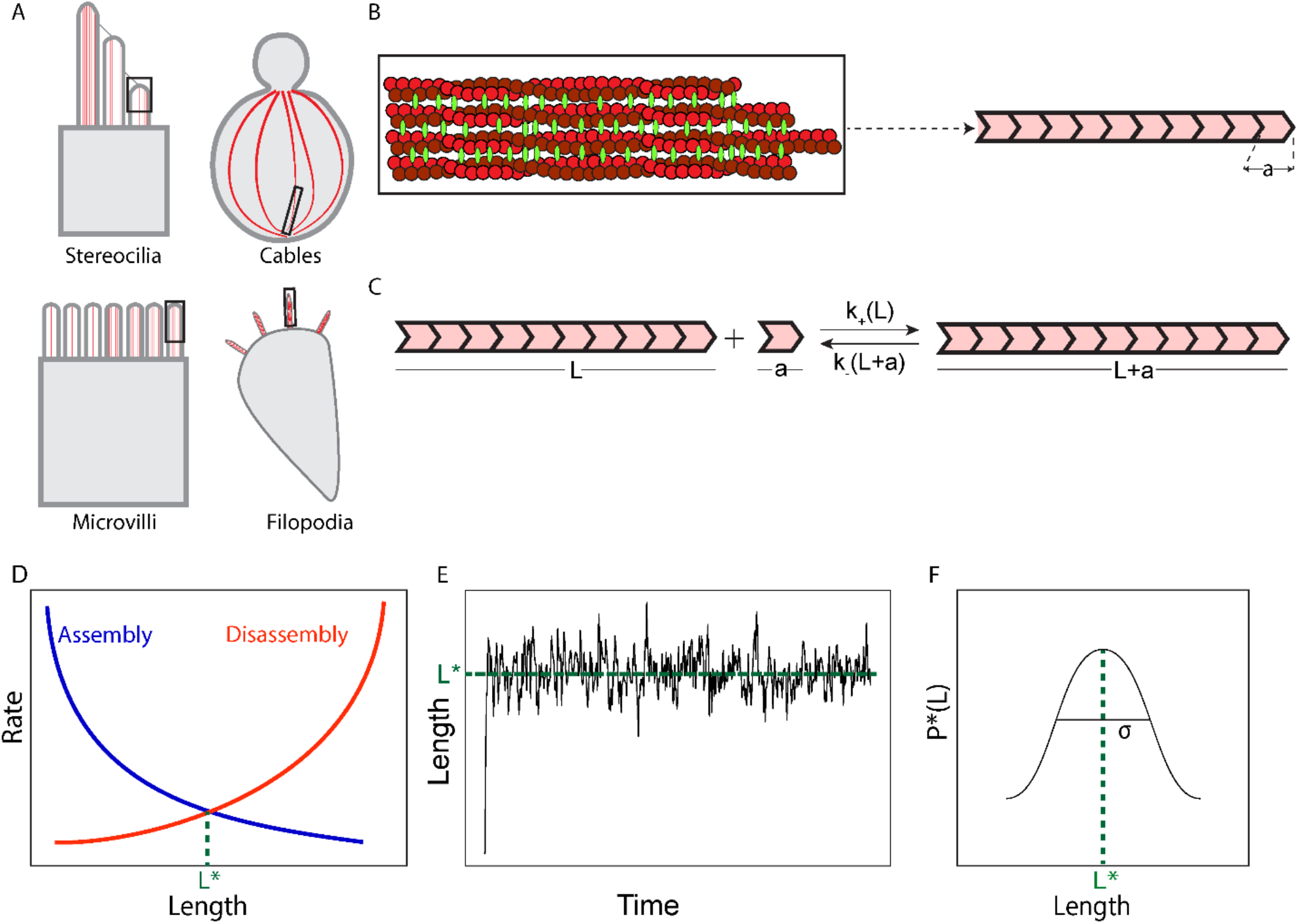
Dynamics of filamentous actin structures in cells: (A) Diverse filamentous actin structures (red) in cells with different functions: (top left and clockwise) stereocilia in hair cells for mechanotransduction, actin cables in budding yeast for intracellular transport, filopodia in motile cells for local environment sensing, and microvilli in epithelial cells for absorption of extracellular chemicals. (B) These filamentous structures all consist of parallel actin filaments (red, zoomed in view of black box) bundled by crosslinking proteins (green). To describe changes of their length over time we model these linear structures as a single polar filament consisting of building blocks (pink chevron tile) of length *a*, in monomer units. (C) In “balance point models” of length control, assembly and disassembly of the polymer, which abstracts the filamentous structure, is described as the stochastic addition and removal of individual building blocks with rate constants *k*_+_(*L*) and *k*_−_(*L*), which can depend on the length of the polymer (*L*). (D) A steady state length, *L*\* is achieved when the two competing rates match each other, at the intersection of the red and blue lines. (E) The length versus time schematic shows a typical output expected from stochastic simulations of the balance point model: the length rapidly grows from *L* = 1 until it reaches a steady state *L*\* and fluctuations around the steady state length are observed. (F) These steady state length fluctuations define the probability distribution function *P*\*(*L*), here shown schematically, which can be characterized by its mean and variance (σ^2^), where σ is the standard deviation of *P*\*(*L*). (The length vesus time graph in (E) was obtained from a model where 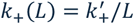 and *k*_−_ (*L*) = *k*_−_; see Figure 2A.)

For individual actin filaments an addition or removal of a single actin protein elongates or shortens the filament by *2.7* nm (accounting for the helical structure of the filament^1^). However, for actin bundles *in vivo*, single events can add or remove larger segments of a filament (consisting of many monomers), leading to a larger change in length. For example, a recent *in vitro* study on depolymerization of actin filaments in the presence of budding yeast CAP (Srv2) and Cofilin (Cof1) found that binding of an individual Srv2 at one end of the filament can induce a depolymerization event that reduces the filament length by *a*_−_ ≈ 270 nm (100 monomers). In fact, the measured depolymerization rate in this case is 300 times greater than in the absence of these two actin binding proteins^24^.

The size of the ‘subunit’ (ranging from a single monomer to many monomers) added to a filamentous structure in a single assembly event (*a*_+_) can also vary depending on the concerted effort of assembly and bundling factors. For example, a recent quantitative *in vivo* study on actin cables in budding yeast raised the possibility that actin filaments of length *a*_+_ ≈ 500 nm are added to the growing cable, which are polymerized by formins and added to the bundle by crosslinking at the bud neck^21^. This type of superstructure for actin cables has also been observed in fission yeast by electron microscopy^25^.

To model the stochastic nature of addition and removal of subunits from a growing filamentous structure, we introduce the rate constant *k*_+_ (*L*) for adding, and *k*_−_ (*L*) the removal of fragments to the effective polymer, which represents the filament (Figure 1C-D). In keeping with the basic tenets of the balance point model, we assume that one or both rate constants, for assembly and disassembly, depend on the length of the polymer, *L*^12^. Here we analyze the situation when the subunit sizes are the same *a*_+_ = *a*_−_ = *a*. We describe the more general result for the case when the two are different in the Supplement, but this does not add anything qualitatively new to the discussion presented here (Supplementary Figure 2).

**Figure 2.**
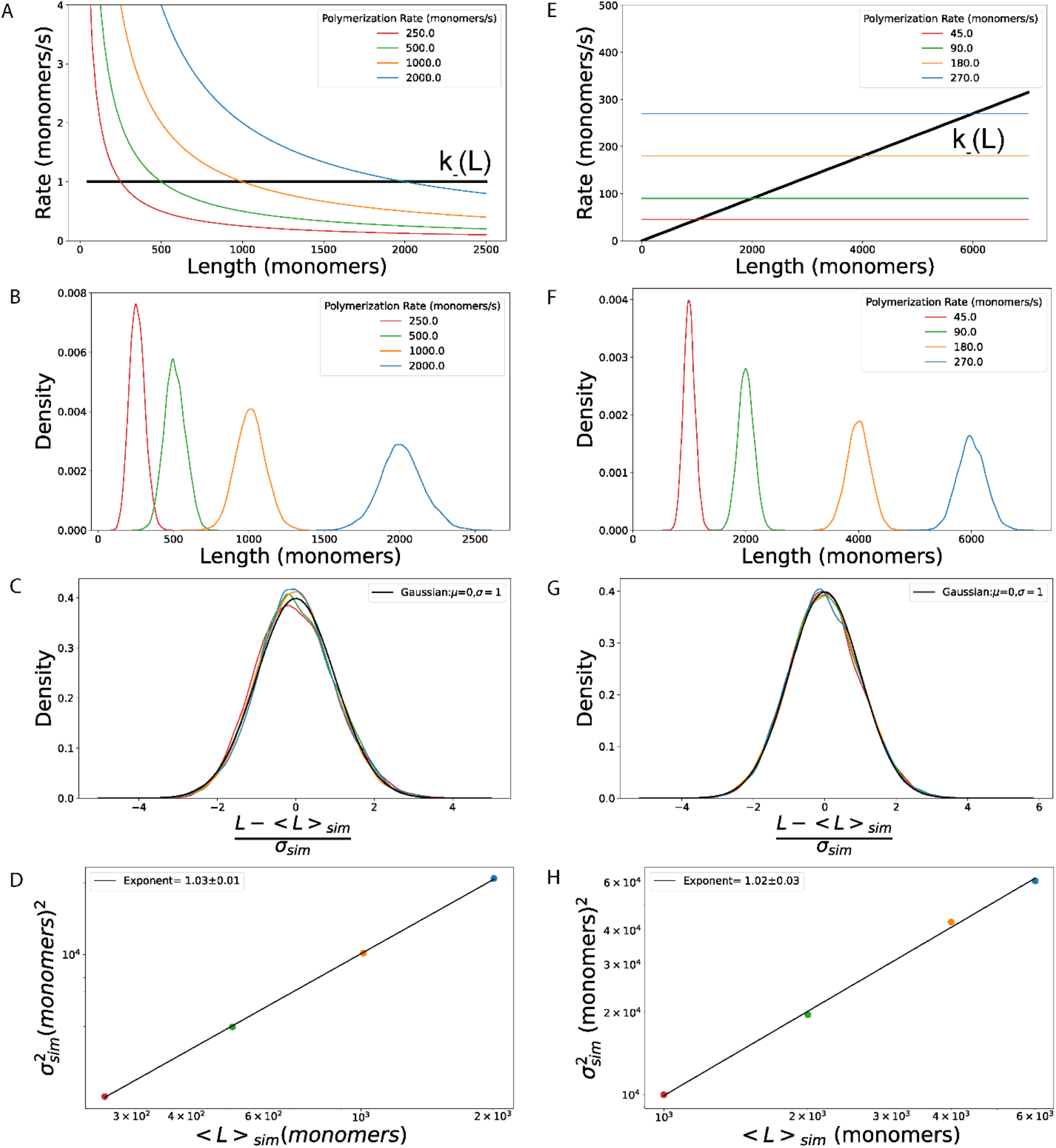
Universal length fluctuations in balance point models: (A-D) Results for a balance point model with a length dependent rate of assembly, *k*_+_ (*L*) = *κ*_+_/*L*, and length-independent disassembly, *k*_−_ (*L*) = *k*_−_. Different steady state lengths, *L*\*, are achieved by tuning the assembly parameter, 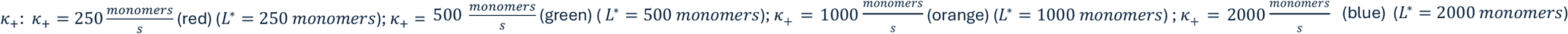. In all simulations the length-independent rate of disassembly, 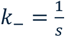. (B) Steady state length distributions from stochastic simulations for different values of *κ*_+_. (C) The length distributions from (B) collapse to a Gaussian distribution (black line) centered around zero with a standard deviation of one, when the lengths are rescaled by the mean and standard deviation of each individual length distribution. (D) The variance of the length distributions scales linearly with the mean length (error bars are standard deviations). (E-H) Results for a balance point model with a length independent rate of assembly, *k*_+_ (*L*) = *k*_+_, and length-dependent disassembly *k*_−_ (*L*) = *κ*_−_*L*. Different steady state lengths, *L*\*, are achieved by tuning the assembly rate 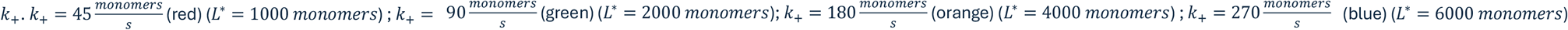. In all simulations the disassembly rate parameter 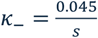 (F) Steady state length distributions for different values of *k*_+_, obtained from stochastic simulations. (G) The length distributions from (F) collapse to a Gaussian distribution (black line) centered around zero with a standard deviation of one, when the lengths are rescaled by the mean and standard deviation of each individual length distribution. (D) The variance of the length distributions scales with the mean length to the power 1.02 ± 0.03. (Error bars are standard deviations). In all the simulations the subunit size is *a* = 10 monomers.

With these assumptions in hand the average length of the polymer (which represents the filamentous structure consisting of many, crosslinked filaments) is described by the dynamical equation:

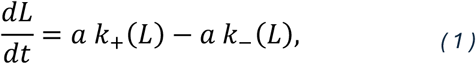

which defines the steady state length of the polymer, *L*\*, as the length for which the two rates are balanced, i.e., *k*_+_ (*L*\*) = *k*_−_ (*L*\*) = *k*\*, where *k*\* is the steady state rate.

Our goal is to compute the length fluctuations in steady state. For this, we introduce *P*(*L, t*), the probability that the polymer has length *L* at time *t*. The time evolution of this probability distribution is described by the chemical master equation:

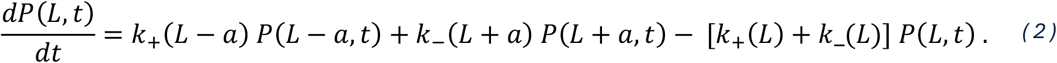

As in any chemical master equation, the right-hand side of Equation 2 describes all the ways in which the probability of the polymer having a length *L* can change in a short time interval. Namely, the polymer can grow from length *L* − a by addition of a subunit with a rate *k*_+_ (*L* − a), or it can shrink from length *L* + a by removal of a subunit with a rate *k*_−_ (*L* + a). These two processes increase the probability of the polymer having length *L*. Alternatively, a polymer of length *L* can grow to length *L* + a with a rate *k*_+_ (*L*) and shrink to a length *L* − a with a rate *k*_−_ (*L*). These two processes decrease the probability of finding the polymer at length *L*. In steady state, the probability distribution is stationary in time, i.e., 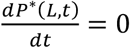, where *P*\*(*L*) ≡ *P*\*(*L, t*) is the time-independent, steady state distribution of polymer lengths.

Since our master equation is a linear Markov chain with no loops, its unique steady state satisfies the detailed balance condition:

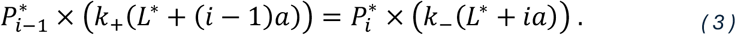

Here the lengths of the polymer are written as *L* = *L*\* + *ia* with *i* taking integer values, and 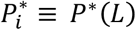; in other words, we use *i* as a dimensionless measure of the polymer length deviation from its steady state value (in units of *a*).

Equation 3, provides a recursive relation for steady state probabilities from which any 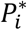 can be computed from the knowledge of 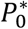, which is the probability that the length is equal to the steady state value, i.e., the deviation of *L* from *L*\* is zero. To simplify this expression, we Taylor expand the rates *k*_+_ (*L*) and *k*_−_ (*L*) around the steady state value of the length (*L*\*):

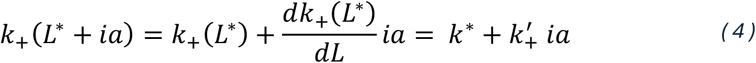

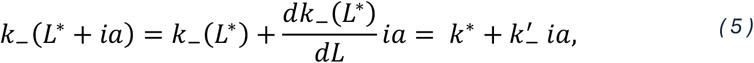

where 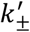 are the derivatives of the assembly and disassembly rates evaluated at the steady state length *L*\*; these are the slopes of the red and blue curves at the point of intersection in Figure 1D. Substituting the Taylor expansions in the condition for detailed balance, Equation 3, results in a recursive formula for the steady state probabilities:

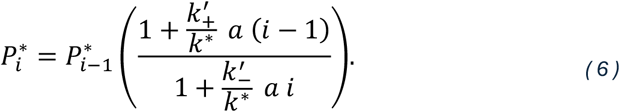

Assuming 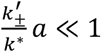, which we show in the Supplement is equivalent to assuming that the steady state polymer length is much bigger than the length of an individual subunit, we can approximate 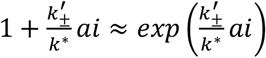. Then, using the normalization condition that requires that the probabilities of all lengths sum up to one, we arrive at a simple formula for the steady state distribution of polymer lengths:

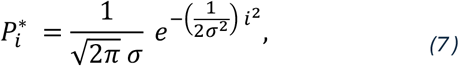

where 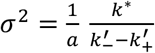 is the variance of the distribution 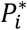. The mean is zero, which just says that, within this approximation the mean and the steady state length are equal.

Therefore, the balance point model predicts that the distribution of lengths at steady state is a Gaussian, irrespective of the functional form of the assembly and disassembly rate constants. Replacing back for the integer variable *i* = (*L* − *L*\*)/*a*, the variance of the polymer length is given by,

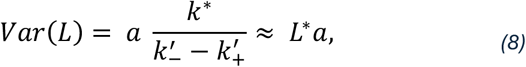

where we approximate the derivatives of the rates in the denominator with the ratio of the steady state rate and the steady state length, i.e., 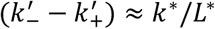 (for details see Supplement). Equation 8 provides us with a key prediction of the balance point model that can be tested in experiments, that the variance of the steady state length distribution scales with the first power of the steady state length.

As we discuss at length in the Supplement, even though Equation 7 approximates the exact result described by Equation 6, the approximation is valid for all polymer lengths that are within 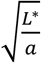 standard deviations (*σ*) of the mean, which, even for the case when the subunit length is only ten times smaller than the steady state length, corresponds to within three *σ*’s of the mean, or 99.7% of the full polymer length distribution. In other words, we expect the Gaussian approximation to be valid for all filament lengths measured in a typical *in vivo* experiment.

To test the two key results of our analysis, namely that the distribution of lengths is Gaussian (Equation 7), and that the variance of the length varies linearly with its steady state value (Equation 8), we performed stochastic simulations of two different balance point models described in the literature. Mathematically speaking, different balance point models correspond to different choices of *k*_+_ (*L*) and *k*_−_ (*L*).

In the first model, we assume that the rate of polymerization falls inversely with the length of the polymer, *k*_+_ (*L*) = κ_+_/*L* while the rate of depolymerization is independent of polymer length, *k*_−_ (*L*) = *k*_−_ (Figure 2A). Such a model has been extensively studied in the context of length regulation of flagella, which are microtubule-based filamentous structures used for swimming, in the single cell algae *Chlamydomonas reinhardtii*^26^. Experiments on these flagella, as well as those on short (< 500*nm*) microvilli in *Xenopus* kidney epithelial cells show a length-dependent assembly rate while the disassembly seems to occur at a constant rate^27^.

To test the two key predictions of our theory, we performed stochastic simulations of this balance point model for different values of the rate parameter κ_+_, which lead to different steady state lengths. Setting the rates of polymerization and depolymerization in steady state to be equal gives the expression for the steady state polymer length: *L*\* = κ_+_/ k_−_; in simulations we varied the steady state length from *L*\* = 25*a* to *L*\* = 200*a*, where *a* is the subunit length.

In Figure 2B we plot the length distributions obtained from the stochastic simulations for different values of κ_”_, while in Figure 2C we demonstrate that they all collapse to the standard Gaussian curve with mean zero and standard deviation of one, once the distributions are plotted as a function of (*L*−< *L* >_*sim*_)/σ_*sim*_, where the mean (< *L* >_*sim*_) and the standard deviation (σ_*sim*_) of the polymer lengths are determined from the simulation. In Figure 2D we show the variance of the length distribution obtained in the simulations plotted against its mean. The straight line on this log-log plot indicates scaling of the variance with the mean to the power 1.03 ± 0.01, consistent with the theoretical prediction of scaling with an exponent of 1. In the Supplement we also compare the results of simulations of this balance point model when k_−_ is varied instead of κ_+_, and once again we find excellent agreement with our theoretical predictions (Supplementary Figures 1A-D).

The second model we explore in stochastic simulations has a length-independent rate of polymerization, *k*_+_ (*L*) = k_+_ and a rate of depolymerization that increases linearly with the length of the polymer, *k*_−_ (*L*) = κ_−_ *L* (Figure 2E). As in the previous case, setting the rate of polymerization and depolymerization at steady state to be equal, we can compute the steady state length, *L*\* =k_+_ / κ_−_. This model has been proposed for length control of microtubules by kinesin motors, such as Kip3 and Kif19 disassembling tubulin monomer from the plus ends of microtubules. The microtubule acts as a landing pad for these motor proteins and longer microtubules have proportionally more of these motors reaching the end of the microtubule per unit time. Therefore, longer microtubules will have a proportionally higher rate of disassembly, i.e., *k*_−_ (*L*) = κ_−_ *L* ^28,29^. A similar model based on formin regulator Smy1 has also been proposed for actin cables in budding yeast^30^. A length-independent rate of polymerization and a length-dependent rate of disassembly has been proposed as the mechanism of length control for microtubule-based flagella in *Giardia*^31^ and for treadmilling filaments in general^32^.

Just as in the numerical exploration of the previous balance point model, here we similarly vary the steady state polymer length by varying the rate of subunit addition k_+_ (Figure 2E). In Figure 2F we show the length distributions obtained for the different values of k_+_ and in Figure 2G we demonstrate that, consistent with one of our key predictions, all the distributions collapse onto the standard Gaussian curve when plotted against (*L*−< *L* >_*sim*_)/σ_*sim*_. Finally, as in the previous model, fitting the variance as a function of the mean to a power law reveals an exponent of 1.02 ± 0.03 (Figure 2H), consistent with our prediction of linear scaling. In the Supplement we also compare the results of simulations of this balance point model when κ_−_ is varied instead of k_+_, and find excellent agreement with our theoretical predictions (Supplementary Figures 1E-H).

In the more general case, when the lengths of subunits that are added (*a*_+_) and removed (*a*_−_) in a single chemical step are different, the theoretical predictions are the same. Namely, the steady state distribution of lengths is Gaussian with a variance that is proportional to the steady state length, *Var*(*L*) ≈ *L*\*(*a*_+_ + a_−_)/2. Details of this calculation can be found in the Supplement, where we also show results of stochastic simulations of a balance point model with *a*_+_ ≠ *a*_−_ that are consistent with these predictions (Supplementary Figure 2).

### Length fluctuations of filamentous actin structures in vivo have a variance that scales quadratically with the mean length

Our calculations show that the balance point models make a quantitative prediction about length fluctuations of filamentous structures, namely that the variance of the length should scale with the mean length to the first power. We test this theoretical prediction in Figure 3 using published experimental results on stereocilia in hair cells, microvilli of epithelial cells, cables in budding yeast, and filopodia in motile cells and we find evidence against it. In addition, we uncover a universal scaling of the variance with the square of the mean length of these filamentous structures.

**Figure 3.**
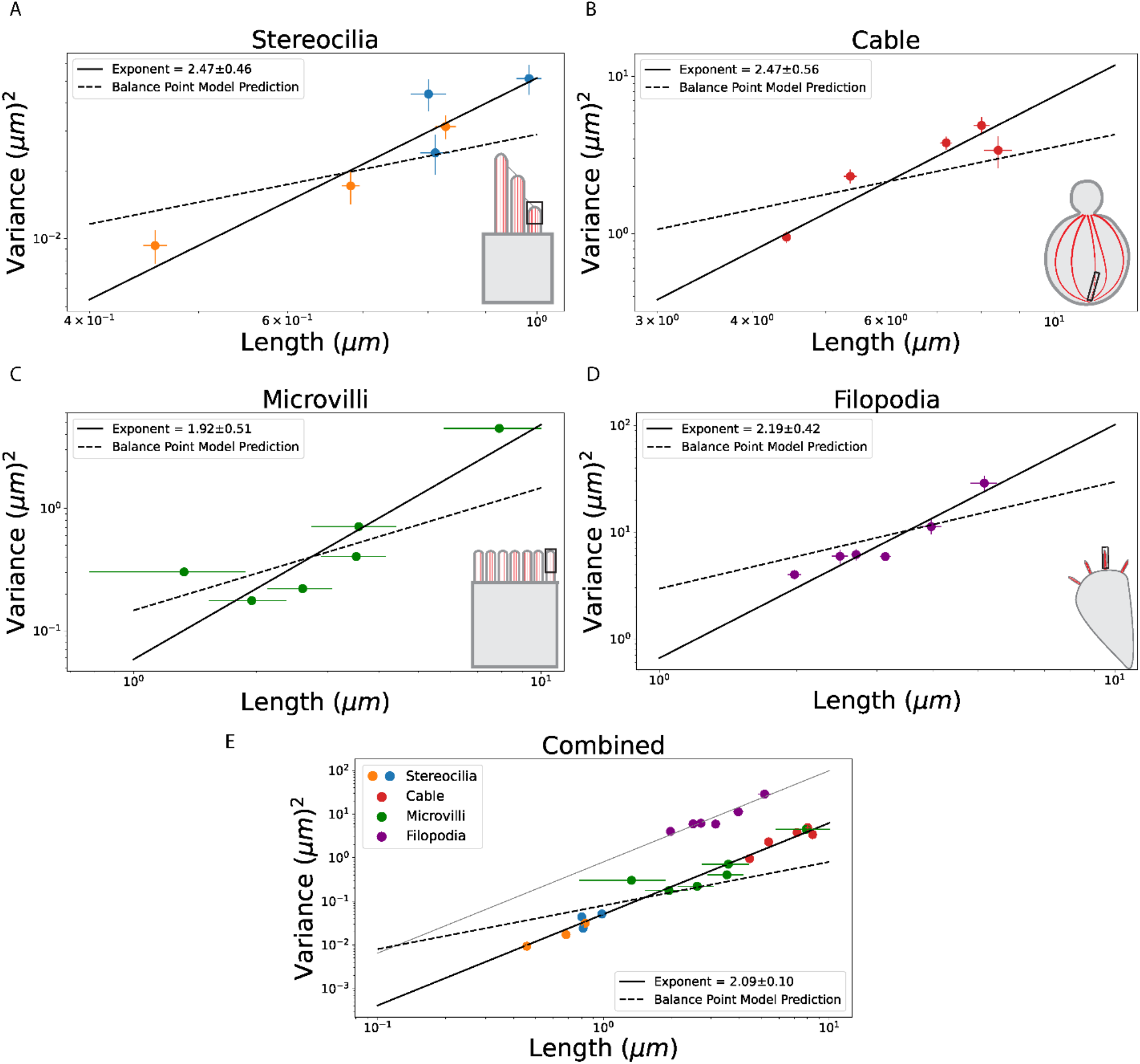
Universal scaling of variance with mean length for filamentous actin structures *in vivo*. (A) Row 3 (shortest) stereocilia in mouse inner (blue) and outer (orange) hair cell bundles show length variation when treated with drugs such as amiloride and benzamil that block the flow of calcium ions in the tip links connecting the shorter rows to the taller ones. The log-log plot of the variance versus the mean of stereocilia lengths has a slope of 2.47 ± 0.46 (*R*^2^ = 0.86). (B) Actin cables (red) in budding yeast have lengths that scale with the length of their mother cells. The log-log plot of the variance versus the mean cable length has a slope of 2.47 ± 0.56 (*R*^2^ = 0.82). (C) Exogenous actin bundling proteins caused length variation in brush border microvilli of epithelial cells (green). The log-log plot of the variance vs mean length of microvilli has a slope of 1.92 ± 0.51 (*R*^2^ = 0.74). (D) Different filopodia lengths are achieved by genetic, chemical, and mechanical perturbations. The log-log plot of the variance vs mean of filopodia lengths have a slope of 2.19 ± 0.42 (*R*^2^ = 0.89). (E) Combined plot of the data for all the different actin structures in (A)-(D). The data for stereocilia, cables, and microvilli fall on a single power-law line with a slope of 2.09 ± 0.10 (*R*^2^ = 0.98). The data for filopodia follows a similar power law scaling as the rest (the gray line is parallel to the black line) but with a larger prefactor. All the errors in the slope are standard errors. Data points show mean ± standard error (orange, blue, red, and purple) except green which is mean ± standard deviation (see Methods for details about error estimates).

It is important to note here that the theory described above deals with length fluctuations that are caused by the stochastic nature of filament assembly and disassembly. It does not consider the effect of possible cell-to-cell variations in rate parameters that define the assembly/disassembly kinetics, that can arise, for example, due to variations in the concentrations of proteins involved in the dynamics of these filaments^33^. The available experimental data is typically from an iso-genetic population of cells and even though contributions from the cell-to-cell variation to length fluctuations is possible, in our analysis we assume that their contributions can be neglected. We examine this assumption in more detail in the Discussion, and argue why we believe it stands to reason, but the final arbiter will be future experiments that measure length fluctuations at the single cell level, like in the recent experiments on microtubule-based flagella^17^.

In Figure 3A, we show data for mouse inner(blue) and outer(orange) hair cell row three stereocilia obtained from electron microscopy images by Ve lez-Ortega et al^18^. The longest lengths in both colors correspond to the wildtype stereocilia and the shorter lengths correspond to stereocilia from cells treated with a mechano-electrical transduction ion current blocking drugs, such as amiloride and benzamil, for 24-hours, which has the effect of reducing their length. This data was used to determine the mean and variance of the stereocilia length which when plotted against each other shows power law scaling with an exponent of 2.47 ± 0.46. This is in contradiction with the linear scaling predicted by the balance point model (shown for comparison as the dashed lines in Figures 3A-E).

In Figure 3B we plot the variance and mean lengths of cables in budding yeast (red)^21^. Genetically perturbed mutant cells with lengths twice that of wild type cells, have actin cables that grow to match the lengths of mother cells, as they do in wild type cells. The wild type haploid, wild type diploid, and mutant cells were grouped by their mother cell lengths into five bins. The variance and mean lengths of cables from each bin was calculated and plotted. We find power law scaling with an exponent of 2.47 ± 0.56, inconsistent with the balance point model prediction.

The variance of microvilli lengths (green) measured in wild type and cells transiently transfected with different levels of actin bundling proteins as a function of their mean lengths are plotted in Figure 3C^19^. The data points correspond to cells: wild type, and expressing 2% espin levels, fimbrin, 10% espin, villin, and 100% espin. Their variance scales with the mean length with a power law of 1.92 ± 0.51, an exponent higher than expected from the balance point model prediction.

Husainy et al. quantified the distribution of filopodial lengths using the rodent cell line Rat2 in wild type cells and after genetic, chemical, and physical perturbation^20^. In Figure 3D, the variance of the filopodia lengths (purple) measured in wild type and variously perturbed cells are plotted against their mean lengths. They show a power-law scaling with a slope of 2.19 ± 0.42, higher than what is predicted by the balance point model.

The data shown in Figures 3A-D are from very different cell types and correspond to functionally very different filamentous actin structures. What all these have in common though, is that they are all bundles of individual actin filaments held together by crosslinking or bundling proteins^2,4,34^. To test whether this geometrical arrangement might be responsible for the observed relation between the mean and variance of the length we plot all the data together in Figure 3E. Remarkably, we see that the data from these disparate cells and very different actin structures all roughly fall on a single curve. The solid line is a fit through stereocilia, microvilli, and cables and corresponds to a power law with an exponent of 2.09 ± 0.10 with an R^2^ value of 0.98. The filopodia data seemingly has the same slope (grey line is the solid line with a higher prefactor) but with variances higher than what is measured for other actin structures.

This shift of filopodia data from the trend defined by the other data sets (Figure 3D-E (purple)) could be due to the way in which filopodial lengths were measured in experiments. The length of the filopodia is defined as the length of the protrusion from the cell membrane to the tip. However, filopodial actin bundles are much longer than the protrusion as they extend significantly into the cytoplasm, as revealed by EM studies^35^. If we assume that the actual length of actin bundles supporting the filopodia is 2-3 times of what it reported, then the filopodia data would align with the other actin structures shown in Figure 3 (see Supplementary Figure 7).

While the scaling observed in the data is of very limited dynamic range, filament lengths range from 0.4*μm* − 10µ*m*, we see a clear violation of the scaling predicted by the balance point model, as indicated by the dashed lines in Figures 3A-E. This is the key conclusion we take away from reanalyzing the experimental data on *in vivo* lengths of filamentous actin structures.

### A “bundled-filament model” predicts the experimentally observed scaling of variance with mean length

To reconcile the discrepancy between theory and experiments described above, we propose a model of length control for parallel actin bundles where we consider the bundles to be made of *N* actin filaments, held together by crosslinking molecules (Figure 4A). We assume that the individual filaments are dynamic, undergoing constant assembly and disassembly, which leads to an exponential distribution of filament lengths in steady state: 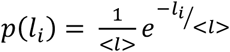, where < *l* > is the average filament length. An exponential distribution of lengths is obtained when the assembly and disassembly rates are independent of length, and disassembly is greater than the assembly rate. In the Supplement, we also consider bundles of filaments whose length is controlled by severing and we arrive at similar conclusions (Supplemental Figure 4).

**Figure 4.**
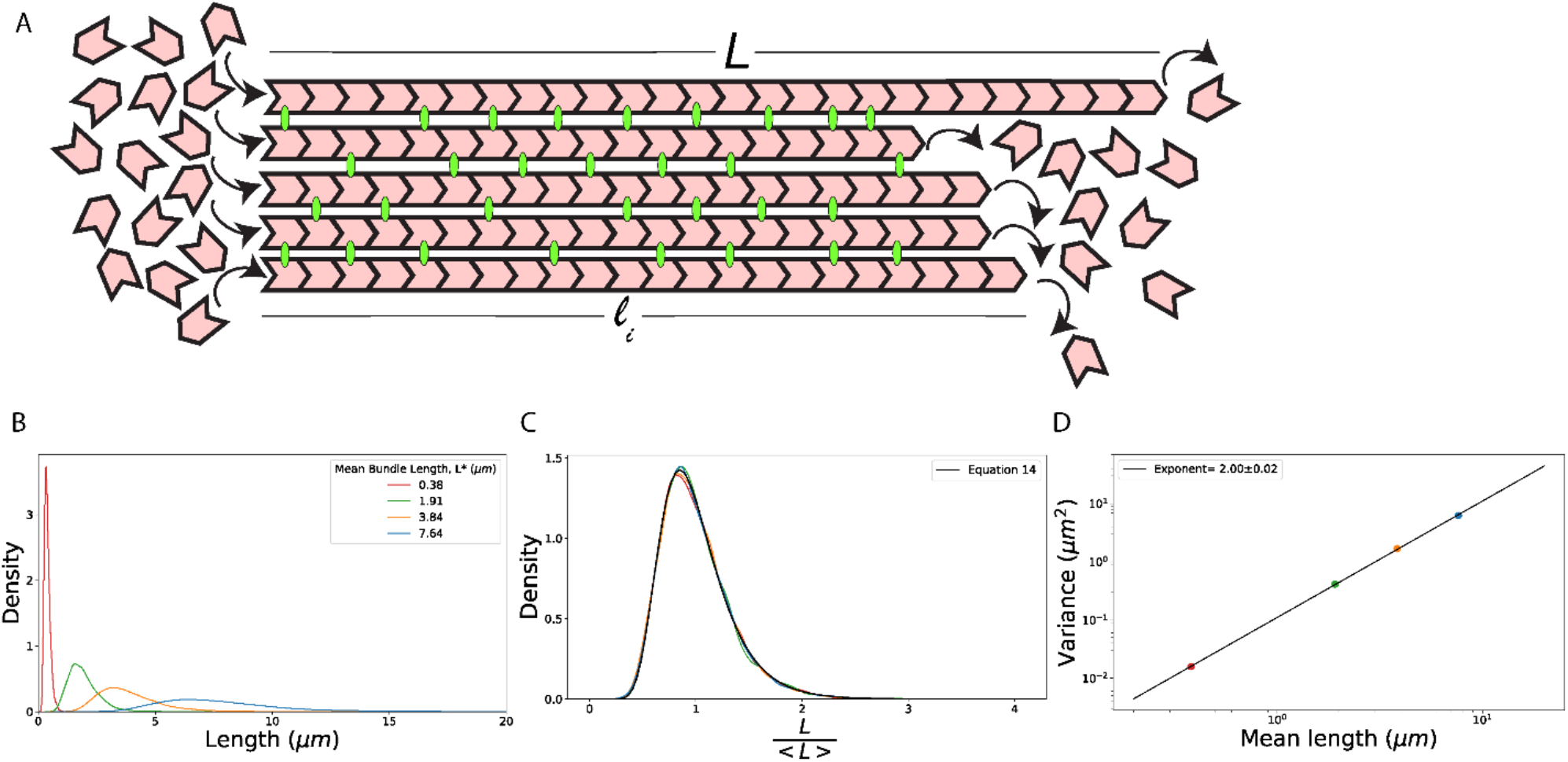
Bundled-filament model of length regulation: (A) We consider parallel actin bundles to be made of *N* actin filaments (red), held together in a bundle by crosslinking proteins (green). Individual filaments have lengths: *l*_*i*=1,…,*N*_, which are assumed to be exponentially distributed. The bundle length is defined as the maximum of all the filament lengths, *L* = *max* {*l*_1_, *l*_2_, …, *l*_*N*_}. (B) Length distribution of a bundle of *N* = 25 filaments obtained from a simulation that generated 10,000 bundles. Individual filaments within a bundle are sampled from an exponential distribution with mean lengths: < *l* > = 0.1 *μm* (red), < *l* > = 0.5 *μm* (green), < *l* > = 1.0 *μm* (orange), and < *l* > = 2.0 *μm* (blue). (C) The distributions of bundle lengths collapse onto a single curve when the lengths are normalized by the mean bundle lengths, which coincides with the theory prediction, Equation 14 (black line). (D) The log-log plot of the variance as a function of the mean length from the distributions showed in (B) gives a power law exponent of 2.00 ± 0.02.

The length of the filament bundle is defined as the length of the longest filament in the bundle, *L* = *max* {*l*_1_, *l*_2_, …, *l*_*N*_}. The probability that any filament in the bundle has a length *l*_*i*_ < *L*, is given by the cumulative distribution function *F*(*L*),

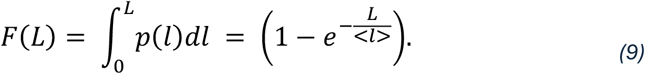

For all the *N* filaments within the bundle to have length less than *L*, the probability is given by *N*^th^ power of this cumulative distribution function,

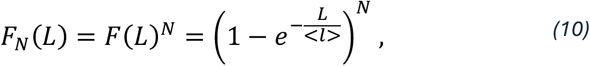

which is simply the mathematical statement that for the bundle to have a length that is smaller than *L* every single filament in the bundle must be shorter than *L*. The probability density function for the longest filament is given by the derivative of the cumulative distribution function,

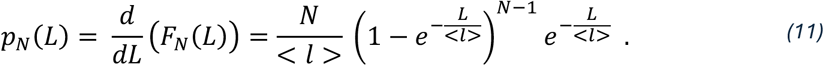

The key result we find is that the distribution of bundle lengths is peaked (Figure 4B) even though each individual filament in the bundle draws its length from an exponential distribution. In other words, by simply bundling individual filaments whose length is not under feedback control, we produce a bundle whose length is controlled.

To characterize the distribution of bundle lengths, from Equation 11 we calculate the average (< *L* >) and variance (*var*(*L*)) of the distribution (for details see Supplement),

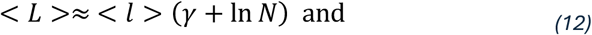

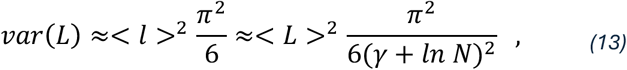

where in Equation 13 we express the variance in terms of the mean length of individual filaments, < *l* > and the mean length of the bundle, < *L* >, *γ* = 0.577 … is the Euler-Mascheroni constant. Equation 12 also implies that the distribution of bundle lengths (Equation 11) when the lengths are normalized by the mean, is only a function of the number of filaments,

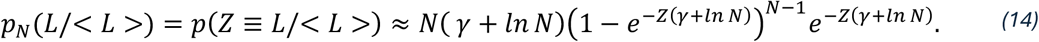

This result is tested numerically in Figure 4C where we show the distributions of the bundle lengths normalized by the mean. All distributions collapse to a single curve given by Equation 14. For large *N* this is the Gumbel distribution used in extreme-value statistics to model the maximum of a number of samples drawn from a common distribution.

Using the bundle model, we derive another key result, that the variance of the bundle length is proportional to the square of its mean length, consistent with the data shown in Figure 3 for all the different filamentous actin structures. Furthermore, we see in Equation 13 that the factor multiplying < *L* >^2^ is logarithmically dependent on the number of filaments (*N*) in the bundle. This could explain why the data for all the different filamentous structures in Figure 3E lie on the same power-law line, even though these structures have different numbers of bundled filaments. Figure 5A illustrates the weak dependence of the average bundle length, < *L* > on the number of filaments in the bundle, *N*, as *N* is varied over 2 orders of magnitude. Also, the variance of the bundle length is independent of the number of filaments when the average filament length, < *l* >, is kept fixed (Figure 5B). In Figure 5C, the theoretical prediction of the variance (Equation 13) is shown for *N* = 10 (top horizontal line) and *N* = 500 (bottom horizontal line), which is the range over which the number of filaments in the different structures vary, as reported for stereocilia^37^, cables^25^, microvilli^38^, and filopodia^39^. Note that the prediction that all the data should fall between these two lines is a sharp, zero-fitting parameter prediction of the bundle model.

**Figure 5.**
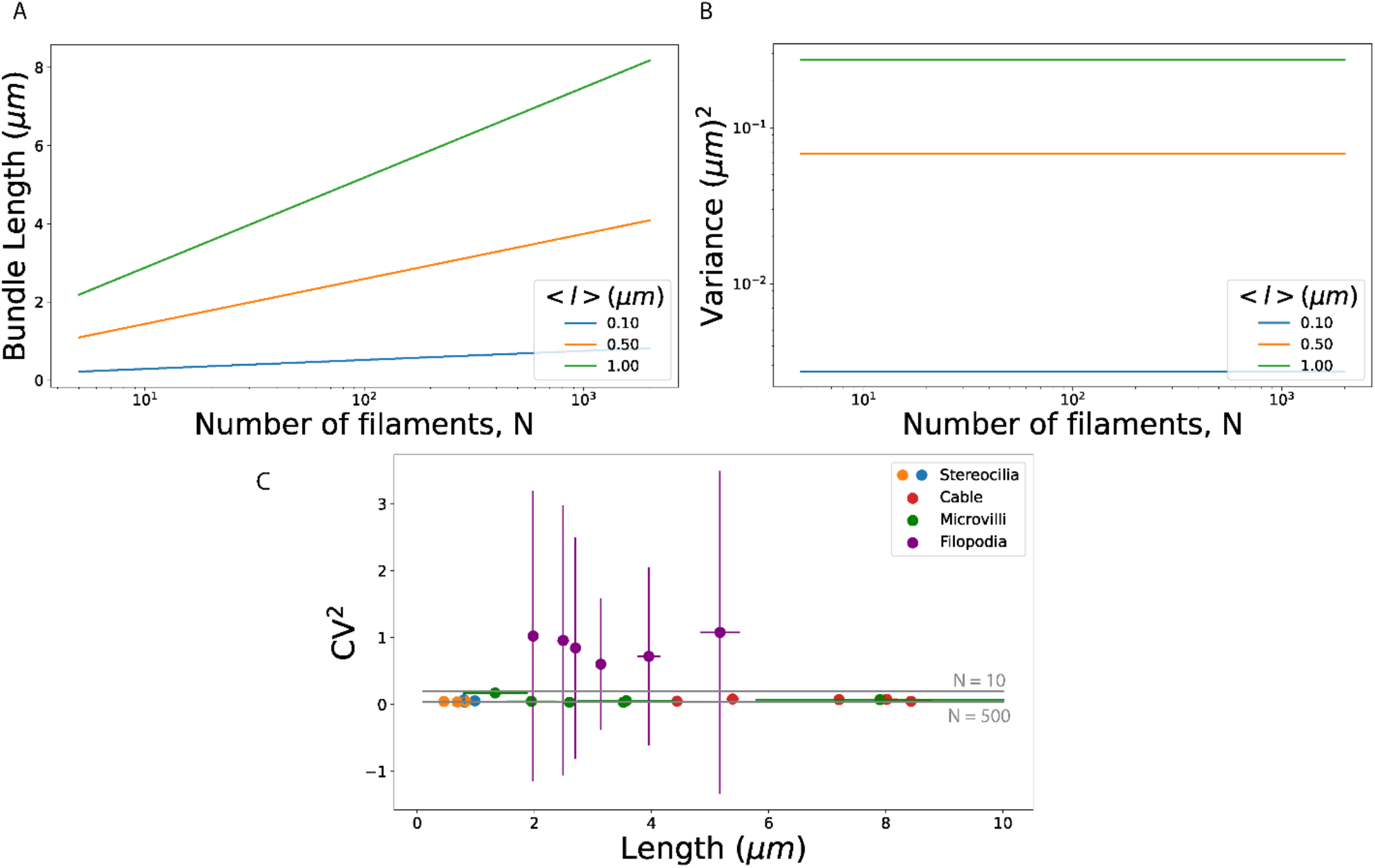
Theoretical predictions of the bundled-filament model: (A) Average length of the bundle, < *L* > as a function of the number of filaments in the bundle, *N* for different average lengths of the exponentially distributed individual filament within the bundle, < *l* >= 0.1 *μm* (blue), < *l* >= 0.5 *μm* (orange), < *l* >= 1 *μm* (green). (B) The variance in bundle lengths is independent of the number of filaments in the bundle when < *l* > is fixed. (C) The square of the coefficient of variation is plotted as a function of the mean length for all the data in Figure 3E, following the same color code. The parallel gray lines are predictions from Equation 13 for different number of filaments in a bundle, (*N*): the top line corresponds to *N* = 10 and the line below corresponds to *N* = 500.

Our model also makes the prediction that the width of the bundle decreases exponentially with distance away from the site of assembly (see Supplement), i.e., the bundle tapers in thickness. This kind of tapering has been observed in stereocilia^40–42^, microvilli^43,44^, bristles^45^, actin bundles in Limulus sperm^46^ and most recently by us for actin cables in budding yeast.

## DISCUSSION

The key result of our paper is that the fluctuations of length predicted by balance point models (Figure 2) are inconsistent with experimental measurements of length for all of the filamentous actin structures we have examined (Figure 3). Instead, we show, that the observed fluctuations can be understood as arising from the bundled nature of these filaments (Figure 4). We show that a model that describes a linear actin structure as a bundle of filaments whose individual lengths are exponentially distributed accounts quantitatively for the experimental data (Figure 5).

### Length fluctuations in balance point models

To compute the length fluctuations in a balance point model we made use of a chemical master equation with length-dependent rates for assembly and/or disassembly, which we solved for to derive the steady state length distribution. We then showed that length fluctuations are well described by a Gaussian distribution (Equation 7) with a variance given by Equation 8.

These results for the length fluctuations in a balance point model can also be derived from the linear noise approximation to the master equation^47^; for details see Supplement. This approximation describes the filament dynamics in steady state as a diffusion process with a diffusion constant for the filament length: *D*_*L*_ = k\**a*^2^. In this approximation, the presence of length-dependent rates leads to an effective restoring force that ‘pushes’ the length back towards the steady state length. This ‘force’ is proportional to the difference between the length and the steady state length of the filament, with a coefficient of restoration 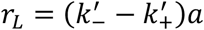. The variance of the length distribution is then simply the ratio *D*_*L*_/*r*_*L*_, which reproduces Equation 8.

The linear noise approximation is expected to be a precise description of the fluctuations away from the steady state regardless of the nature of the assembly process. It applies if the assembly dynamics can be described as the addition and removal of subunits whose size is much smaller than the steady state size of the entire structure. Therefore, we expect the linear scaling of the variance of the length with the steady state length to be a general feature of any model of assembly with these properties.

Filament severing leads to an interesting violation of one of the key assumptions of the linear noise approximation. When a filament is severed, the sizes of fragments released are proportional to the length of the filament before severing, a clear violation of the assumption that the pieces being removed are small. Indeed, the steady state distribution of lengths of a filament undergoing severing, while simultaneously undergoing subunit assembly, is not Gaussian but is described by the Rayleigh distribution (see Supplementary Figure 3B-C). Notably, the variance of this distribution scales with the square of the mean length (Supplementary Figure 3D). Nevertheless, we do not consider severing to be a good model for the length regulation of the filamentous actin structures studied here (Figure 1A), since it is unlikely that the whole bundle would be cut in one chemical step (Supplementary Figure 3A), as this would require severing of all the individual filaments in the bundle at the same location, and at the same time.

### Length regulation and length fluctuations of filament bundles

Many cellular actin structures contain filaments organized into bundles. Here, we have shown the mere organization of filaments into a bundled arrangement endows the resulting structure with a peaked distribution of bundle lengths, i.e., length control. The mean and the standard deviation of the bundle length, defined as the length of the longest filament, is set by the number of filaments and the average length of the filaments that comprise the bundle (Equation 12-13). Therefore, proteins that affect assembly or disassembly of the filaments in the bundle can influence the bundle length simply by changing the mean filament length within the bundle. Interestingly, our model also predicts that the bundle width should fall off as an exponential function of the distance from the growing end, with a decay constant (which describes bundle width tapering) given by the mean filament length. Therefore, a molecular perturbation that changes the mean filament length is predicted to change both the bundle length and its width profile in a proportional manner, i.e., doubling the length of the bundle doubles the distance over which its width tapers. This type of scaling of the width profile of a bundle with its length was recently observed in experiments on actin cables in budding yeast.

To compute the dependence of the variance on the mean bundle length we simply need to know the length distribution of individual filaments. In the supplement, we consider a bundle model where individual filaments within the bundle undergo polymerization by monomer addition at one end and severing along their lengths (Supplementary Figure 4A-D). We found striking similarities between the predictions of this model and the experimental data (Supplementary Figure 5C).

Importantly, our model works for either a bundle comprised of filaments with an exponential distribution of lengths, or for a bundle comprised of filaments whose lengths are controlled by severing and follow a Rayleigh distribution. Based on the available data, we cannot distinguish between these two possibilities. However, in the future it may be possible to distinguish between these possibilities using electron-microscopy to directly measure filament lengths within bundles found in cells.

In the Supplement, we also consider a Gaussian distribution for individual filaments, as one would expect in the presence of a balance point mechanism controlling their lengths (Supplementary Figure 6A-D). In this case the variance of the bundle length scales as the first power of the mean, contrary to experimental data shown in Figure 3.

### Cell to cell variability and length fluctuations

A key assumption of our analysis is that within a cell population the measured variability of filament lengths serves as a good proxy for the length fluctuations that would be observed in a single cell over time. For example, this assumption would be false if there existed significant cell to cell variability in a factor that influences the filament assembly/disassembly processes and therefore filament length.

To investigate this possibility further we consider the balance point model with a constant polymerization rate *k*_+_ and a length-dependent depolymerization rate *k*_−_ (*L*) = *κ*_−_*L* (see Figure 2E), now with an added assumption that the polymerization rate varies from cell to cell. In this case, the variance of the filament lengths has a contribution both from the stochastic process of adding and removing subunits (as analyzed in Figure 2E-H) and from the cell-to-cell variability of *k*_+_. The total variance of the length fluctuations can be computed from the law of total variance (see Supplement for details):

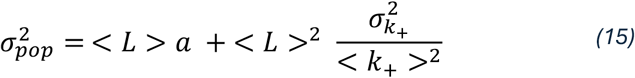

where < *k*_+_ > and 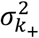 are the population mean and variance of the polymerization rate.

For this model to explain the experimental data two things must be true. First, variance of the cell-to-cell distribution of the rate parameter *k*_+_ must scale with the square of the mean 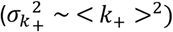. Second, the ratio of these two values would have to be the same for microvilli, cables, filopodia, and stereocilia (see Figure 5C). It seems unlikely that both would be true for the following reasons.

First, it has been demonstrated experimentally that all proteins expressed in cells, including key actin regulatory proteins that control bundle formation, exhibit cell to cell variability. Contrary to the first requirement above, measurements of the variance, over a wide range of conditions, reveal scaling with the first power of the mean (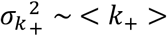)^48^. Second, given the differences in molecular mechanisms leading to the formation of microvilli, cables, filopodia, and stereocilia, we consider it improbable that they all end up producing an effective rate constant (e.g., *k*_+_) with the same coefficient of variation 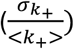 across a population.

In conclusion, by reanalyzing published data, we find that the length fluctuations among diverse bundled actin structures in cells have a remarkable universal feature, that the variance of the bundle length scales with the square of its mean. Furthermore, we show theoretically that this finding is contrary to the predictions of balance-point models for length control, yet in excellent agreement with models that consider the geometry of a bundle. Finally, our results emphasize the importance of measuring the size fluctuations of cytoskeletal structures found in cells, as they can effectively rule out different models of size control.

## METHODS

### Stochastic simulations of balance point models

We simulated the stochastic trajectories of polymer length using the Gillespie algorithm^49,50^ with assembly and disassembly rates that define the balance point models, as described in Figures 2A and 2E. Simulations start with a polymer of length 1, and at each simulation step a subunit is added or removed with probabilities set by the two rates. The time elapsed between consecutive steps are drawn from an exponential distribution with a rate constant given by the sum of the assembly and disassembly rates at a given polymer length. Lengths of the polymer are recorded once the system is in steady state, and these are used to compute the mean and the variance of the steady state distribution (Figures 2D and 2H), as well as the distribution itself (Figures 2B and 2F).

### Bootstrapping method for estimating errors from length data

In making Figure 4 we used published data which reported measured filament lengths. From these we computed the mean and variance and used bootstrapping to estimate the error bars for these two quantities. In each case we resampled the data with replacement, 50,000 times. Standard error of the mean and variance were computed from the resampled measures of mean and variance^51^.

For the stereocilia data the number of filaments measured under different experimental conditions was 53 to 108, while the number of filaments in the filopodia data set ranged from 788 to 1690. The number of filaments in each of the five length bins in the actin stereocilia data ranged from 30 to 300.

### Bundled filament model simulation

To generate a configuration of a bundle we assumed that it consists of 25 filaments whose lengths were sampled from an exponential distribution (Figure 4). For each bundle of 25 filaments, length of the bundle is defined as the length of the longest filament within the bundle. 10,000 such bundles were generated for each case, for which we computed the mean and the variance (Figure 4D), as well as the bundle length distribution (Figure 4B).

## ACKNOWLEDGEMENTS

We thank Ariel Amir, Lishibanya Mohapatra, Alison Wirshing, and Thomas Fai for useful discussions about actin length control and for reading and providing comments on the manuscript. This research was supported by the Brandeis University National Science Foundation (NSF) Materials Research Science and Engineering Center (MRSEC) Bioinspired Soft Materials DMR-2011846, by a grant from the NIH (R35 GM134895) to B.L.G., and the Simons Foundation to J.K.

