## Supplementary text for "Universal length fluctuations of actin structures found in cells"

**SUPPLEMENTARY MATERIAL**

**Calculation of the balance point model length distribution:**

Here we present details of the analytic calculation of the probability distribution of lengths for a polymer whose length is controlled by a balance-point model, with a particular focus on the approximations that lead to the Gaussian form of the distribution.

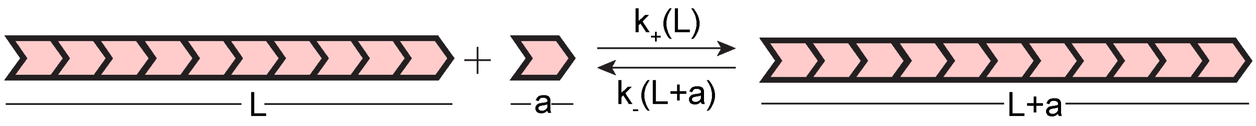


The balance point model is defined by the two length-dependent rates: $k_+(L)$, the rate of polymerization, and $k_-(L)$ the rate of depolymerization. Polymerization corresponds to the addition of a subunit of length $a$, while depolymerization is the removal of a subunit of length $a$ from the filament, as shown in the figure above. The length of the filament changes with time according to the rate equation:

$$\frac{dL}{dt} = k_+(L) - k_-(L)$$

*S1*

The steady state length $L^*$ is the length at which the two rates on the right-hand side of Equation S1 equal each other. The unique steady state satisfies the detailed balance condition,

$$P^*(L-a)\,k_+(L-a) = P^*(L)k_-(L)$$

*S2*

where $P^*(L)$ is the steady state length distribution.

We can write the polymer length $L = L^* + ia$ , where $i$ takes on integer values and counts the number of subunits by which the length deviates from its steady state value. Taylor expanding the rate of polymerization and depolymerization around the steady state length:

$$k_+(L^* + ia) = k_+(L^*) + \frac{dk_+(L^*)}{dL} ia$$

*S3*

$$k_-(L^* + ia) = k_-(L^*) + \frac{dk_-(L^*)}{dL} ia$$

*S4*

leads to a recursive formula for the steady state distribution, $P_i^* \equiv P^*(L)$:

$$P_i^* = P_{i-1}^* \left( \frac{1 + \frac{k'_+}{k^*} a(i-1)}{1 + \frac{k'_-}{k^*} a\, i} \right).$$

S5

To simplify the algebra here we introduced new variables:

$$k^* \equiv k_+(L^*) = k_-(L^*)$$

S6

$$k'_+ \equiv \frac{dk_+(L^*)}{dL}$$

S7

$$k'_- \equiv \frac{dk_-(L^*)}{dL}$$

S8

for the steady state polymerization and depolymerization rates (which are equal; Equation S6), as well as for the derivatives of these two rates evaluated in steady state (Equations S7 and S8).

The recursive formula, Equation S5, can be recast in simpler form assuming $\frac{k'_\pm\, a}{k^*} \ll 1$ (the justification for this approximation is given in the next section), and using the approximate formula $e^x \approx 1 + x$ for $x \ll 1$:

$$P_i^* \approx P_{i-1}^* e^{-\frac{k'_- - k'_+}{k^*} a\, i} .$$

S9

To obtain Equation S9 we have also made the approximation $i - 1 \approx i$ in anticipation that the typical value of $i$ in steady state is much greater than one. The recursive formula in Equation S9 can be solved in terms of the probability that the length is equal to the steady state length ($P_0$):

$$P_i^* \approx P_0^* \prod_{j=1}^{i} e^{-\frac{k'_- - k'_+}{k^*} a\, j}$$

S10

$$P_i^* \approx P_0^*\, e^{-\frac{k'_- - k'_+}{k^*} a \frac{i^2}{2}}$$

S11

This is a Gaussian distribution with zero mean and a variance given by,

$$Var(i) = \frac{1}{a} \frac{k^*}{k'_- - k'_+}$$

S12

We can obtain a simple, approximate expression for the variance by approximating the expression in the denominator of Equation S12 with $\frac{k^*}{L^*}$, i.e., by the ratio of the characteristic assembly/disassembly rate and the characteristics length in steady sate. This approximation amount to assuming that the length dependent rate constants appreciably vary over length scales of order $L^*$

To illustrate this approximation for the expression $k'_- - k'_+$ in steady state, consider the example of the length dependent polymerization rate $k_+ = \kappa_+/L$ and a constant depolymerization rate $k_-$. In this case $k'_- - k'_+ = \frac{\kappa_+}{L^{*2}}$ which at steady state length $L^* = \kappa_+/k_-$ evaluates to $k^*/L^*$, since in this case $k^* = k_- = \frac{\kappa_+}{L^*}$. In a more general setting, we can define the "feedback function"

$$f(L) \equiv k_-(L) - k_+(L)$$

S13

which in dimensionless form can be written as

$$f(L) = k^* F(\frac{L}{L^*})$$

S14

where we assume that the relevant rate scale is $k^*$ and the relevant length scale is $L^*$. The steady state condition is $F(1) = 0$. The derivative of the feedback function evaluated at the steady state length is $f'(L^*) = \frac{k^*}{L^*} F'(1)$. Assuming $F'(1)$ is of order one leads to the scaling described above.

The approximation $k'_- - k'_+ \approx k^*/L^*$ leads to an estimate for the variance in Equation S12,

$$Var(i) \approx \frac{L^*}{a}$$

S15

Changing variables from $i$ back to $L$:

$$Var(L = L^* + ia) = a^2\, Var(i) \approx L^* a$$

S16

gives the key result, that for any balance point model the variance of the polymer length will vary with the first power of the steady state length, and is proportional to the subunit length $a$. Also, from Equation S11 we conclude that the mean length is approximately $L^*$, since the mean deviation $< i > = 0$. Therefore, we predict, for any balance point model, that the variance of the length distribution is proportional to the mean length in steady state.

In the main text we tested the validity of the predicted scaling (Equation S16) using stochastic simulations (Figure 2) for a particular choice of length dependent rates of polymerization and depolymerization. For completeness, in Supplementary figures 1A-D and 1E-H, we show analogous results for the same models used in the main text, but here we vary the rate parameters which were kept constant in the main text. We once again confirm the prediction for the distribution of lengths being Gaussian (Equation S11) and the scaling relation between the variance and the mean (or steady state) length (Equation S16).

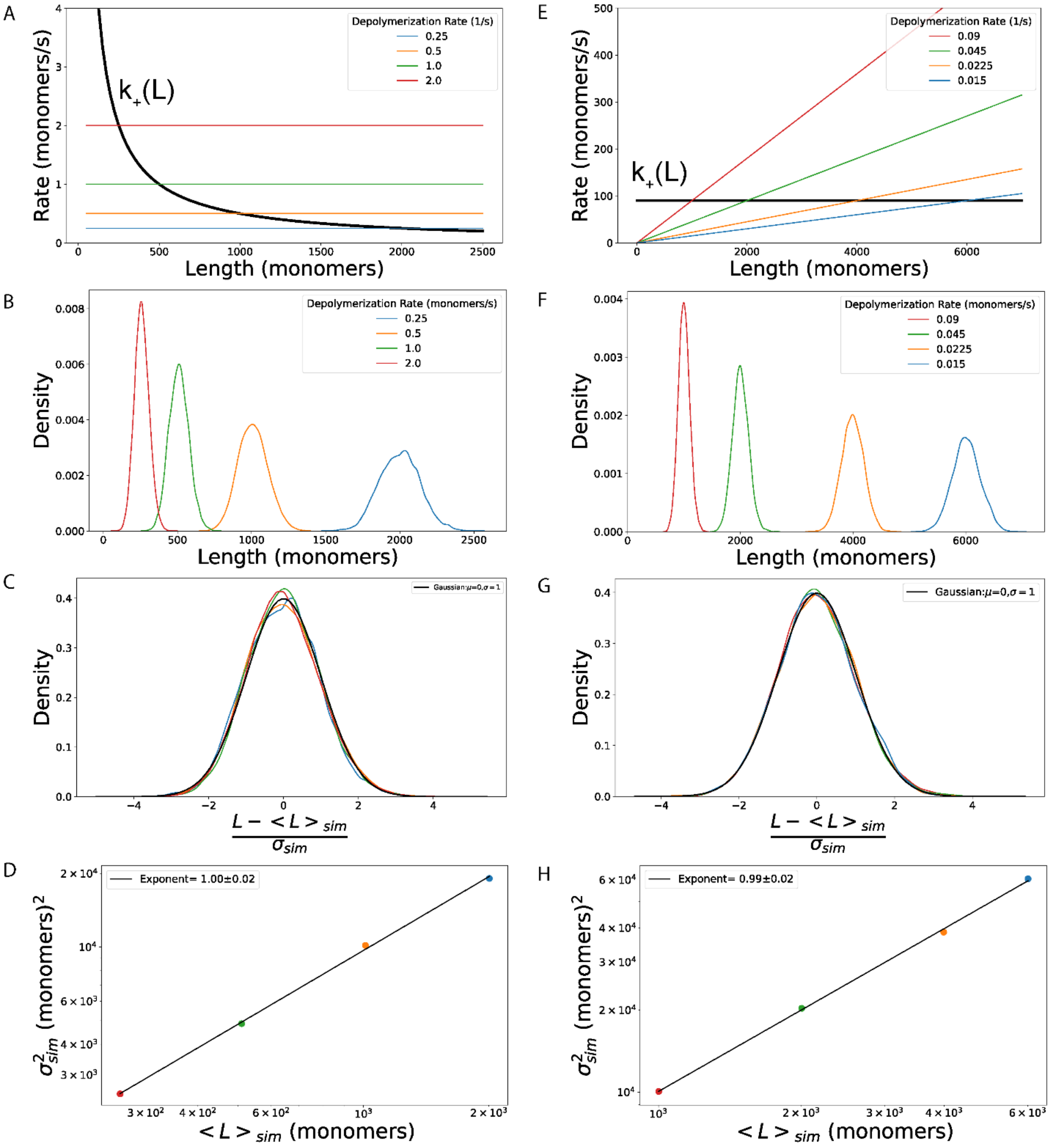


*Supplementary Figure 1: (A-D)* Results for a balance point model with a length dependent rate of polymerization, $k_+(L) = \kappa_+/L$, and length-independent depolymerization rate, $k_-(L) = k_-$. Different steady state lengths, $L^*$, are achieved by tuning the disassembly parameter, $k_-$. $k_- = 2\ s^{-1}$ (red) ($L^* = 250\ monomers$); $k_- = 1\ s^{-1}$ (green) ($L^* = 500\ monomers$); $k_- = 0.5\ s^{-1}$ (orange) ($L^* = 1000\ monomers$) ; $k_- = 0.25\ s^{-1}$ (blue) ($L^* = 2000\ monomers$). In all simulations the length-dependent rate of assembly, $k_+ = 500\ monomers\ s^{-1}$. (B) Steady state length distributions from stochastic simulations for different values of $\kappa_+$. (C) The length distributions from (B) collapse to a Gaussian distribution centered around zero with a standard deviation of one, when the lengths are rescaled by the mean and standard deviation of each individual length distribution. (D) The variance of the length distributions scales linearly with the mean length (error bars are standard deviations). (E-H) Results for a balance point model with a length independent rate of assembly, $k_+(L) = k_+$, and length-dependent assembly $k_-(L) = \kappa_- L$. Different steady state lengths, $L^*$ , are achieved by tuning the disassembly parameter, $\kappa_-$. $\kappa_- = 0.09\ monomers\ s^{-1}$ (red) ($L^* = 1000\ monomers$) ; $\kappa_- = 0.045\ monomers\ s^{-1}$ (green) ($L^* = 2000\ monomers$); $\kappa_- =$

$0.0225\ monomers\ s^{-1}$ (orange) ($L^* = 4000\ monomers$); $\kappa_- = 0.015\ monomers\ s^{-1}$ (blue) ($L^* = 6000\ monomers$). In all simulations rate of polymerization, $k_+ = 500\ s^{-1}$. (F) Steady state length distributions for different values of $k_+$, obtained from stochastic simulations (G) The length distributions from (F) collapse to a Gaussian distribution centered around zero with a standard deviation of one when the lengths are rescaled by the mean and standard deviation of each individual length distribution. (D) The variance of the length distributions scales linearly with the mean length (error bars are standard deviations). In all the simulations the subunit size is $a = 10$ monomers.

**Gaussian approximation of the steady state length distribution:**

Here we show that the Gaussian approximation (Equation S11) to the steady state length distribution of a general balance point model (Equation S1) is valid for all lengths that are within $\sqrt{\frac{L^*}{a}} \gg 1$ standard deviations of the mean length. The key approximation employed in Equation S5 can be rewritten as:

$$f(n) = \prod_{i=1}^{n} \frac{1}{1+\alpha i} \approx e^{-\frac{\alpha}{2}n^2}$$

*S17*

with $\alpha$ given by:

$$\alpha = \left(\frac{k'_- - k'_+}{k^*}\right) a \approx \left(\frac{k^*}{L^*}\right)\left(\frac{a}{k^*}\right) = \frac{a}{L^*}$$

*S18*

For subunit length ($a$) that is nanometer in scale and a polymer that is microns in length, we expect $\alpha \sim 0.001$. To estimate the error made in Equation S17 we use the Taylor expansion:

$$\frac{1}{1+\alpha i} = e^{-\ln(1+\alpha i)} = e^{-\alpha i + \frac{(\alpha i)^2}{2} - \frac{(\alpha i)^3}{3} + \cdots} \approx e^{-\alpha i + \frac{(\alpha i)^2}{2}}$$

*S19*

The first term in the above expression gives the approximation in Equation S17, the second term in the above equation enables us to estimate the size of the error in making this approximation:

$$f(n) \approx \prod_{i=1}^{n} e^{-\alpha i + \frac{(\alpha i)^2}{2}} = e^{-\alpha \sum_{i=1}^{n} i + \frac{\alpha^2}{2}\sum_{i=1}^{n} i^2}$$

$$\approx e^{-\alpha \frac{n^2}{2}\left(1 + \alpha \frac{n}{3}\right)}$$

*S20*

Therefore, for the approximate formula in Equation S17 to be valid we need $\alpha n \ll 1$ in which case we can safely ignore $\alpha \frac{n}{3}$ when compared to 1 in the expression $(1 + \alpha \frac{n}{3})$ appearing in Equation S20.

The formula in Equation S17 is a Gaussian with mean zero standard deviation, $\sigma = \frac{1}{\sqrt{\alpha}}$ . Therefore, the condition $\alpha n \ll 1$ is satisfied as long as $n$ is within $\frac{1}{\sqrt{\alpha}} \gg 1$ standard deviations away from the mean of

$f(n)$. For small $\alpha$ this will be practically the whole support of the function $f(n)$, i.e., values of $n$ for which the condition $\alpha n \ll 1$does not hold are extremely unlikely, as they would be many standard deviations off the mean.

If we restore the dimensional length, we conclude that the Gaussian approximation to the length distribution (Equation 7 in the main text), is a good approximation of the exact solution of the Master equation (Equation 3 in the main text) for all polymer lengths that are within $\sqrt{\frac{L^*}{a}}$ standard deviations ($\sigma \approx (L^* a)^{1/2}$) of the mean. Even for the case when the subunit length is only ten times smaller than the steady state length (i.e., $\frac{L^*}{a} = 10$), this corresponds to three $\sigma$'s off the mean, or 99.7% of the full polymer length distribution. (In reality, we expect the ratio $L^*/a$ to be closer to a 1000.) In other words, if the polymer length is controlled by a balance point mechanism, we do not expect experimental measurements of length, achievable by current methods, to ever detect deviations from a Gaussian distribution.

**Balance point model with unequal subunit segment lengths ($\boldsymbol{a_+ \neq a_-}$):**

In the model described in the main text we have assumed that the subunits added and removed from the filament are of equal length. Here we consider a more general case, when the length of subunits added and removed are unequal, $a_+ \neq a_-$. We also introduce a Langevin equation approach to computing the steady state length fluctuations, which gives the same formula for the variance as we have derived from the Master equation.

When $a_+ \neq a_-$ we can treat the addition and removal of subunits as a random walk (of the polymer length, $L(t)$) with unequal step sizes in the forward and reverse directions, with rates of stepping given by $k_+(L)$ and $k_-(L)$, respectively. To analyze the steady state fluctuations in length we consider this random walk in $L$ around the steady state length $L^*$. In this case we can describe the dynamics of the polymer length by the stochastic Langevin equation[1]:

$$\frac{dL}{dt} = k_+(L) - k_-(L) + \eta(t) \tag{S21}$$

where $\eta(t)$ is a Gaussian random variable, uncorrelated in time, with zero mean and variance, $< \eta(t)\eta(t') > = 2D\delta(t - t')$; it describes the random excursions of the length away from the steady state value.

To compute the "diffusion constant", $D$, we consider the length fluctuations corresponding to adding or removing a single subunit when the polymer length is at its steady state value. In this case the displacement (change in length of the filament), $< \Delta L >_1$, in a small-time interval $\Delta t$ given by:

$$< \Delta L >_1 = k_+(L^*)\, \Delta t\, a_+ \; - \; k_-(L^*)\, \Delta t\, a_- \; = \; 0 \tag{S22}$$

since in steady state:

$$k_+(L^*)\, a_+ \;=\; k_-(L^*)\, a_-$$

S23

The displacement variance after one step is given by

$$< \Delta L^2 >_1 - < \Delta L >_1^2 = k_+^* \,\Delta t\;\; a_+{}^2 \;+\; k_-^* \,\Delta t\, a_-{}^2$$

S24

where we have defined, $k_+^* \equiv k_+(L^*)$and $k_-^* \equiv k_-(L^*)$. Comparing this expression to the variance formula from the Langevin equation for one step away from the steady state, which is given by the canonical diffusion formula $< \Delta L^2 >_1 = \; 2D\Delta t$, we arrive at the formula for the diffusion constant,

$$D \;=\; \frac{1}{2}(k_+^* \, a_+{}^2 \;+\; k_-^* \, a_-^2\;)$$

S25

If we define a step size ratio (or bias), $n = a_-/\, a_+$, and make use of Equation S23, we can rewrite the expression in Equation S25 as:

$$D = \frac{1}{2} k^* a^2 (1+n)$$

S26

where $k^* \equiv k_+(L^*)$ and $a \equiv a_+$.

From the Langevin equation we can now compute the variance of the steady sate length fluctuations as the ratio of the diffusion constant and the "restoring force" for the polymer length:

$$Var(L) = \frac{D}{\frac{d}{dL}(k_+(L)a_+ - k_-(L)a_-)|_{L=L*}}$$

S27

This is the same equation that we derived above using the Master equation. As we did previously, we can make an order of magnitude estimate $\frac{d}{dL}(k_+(L)a_+ - k_-(L)\,a_-)|_{L=L*} \approx \frac{k*\,a}{L*}$, which assumes that both rates vary significantly only on length scales set by the steady state length. Replacing this result and Equation S26 into Equation S27 leads to an estimate of the variance:

$$Var(L) \approx \frac{1}{2} L^* a(1+n)$$

S28

Note that this result reduces to Equation 8 (in the main text) for $n = 1$ when the two lengths of the two subunits are equal. This result for the variance holds under assuming that the length of the subunits that are added and removed are much smaller than the length of the polymer, i.e., $a_+,\ a_- << L^*$.

In Supplementary Figure 2 we show results for stochastic simulation for the models and parameters used in Figures 2A and 2E, now with unequal subunit lengths, $a_+ = 200$ monomers and $a_- = 100$ monomers ($n = 2$). The results of simulations are in agreement with the above calculations; in particular we test the prediction that the length distribution is Gaussian with the variance that scales linearly with the mean length.

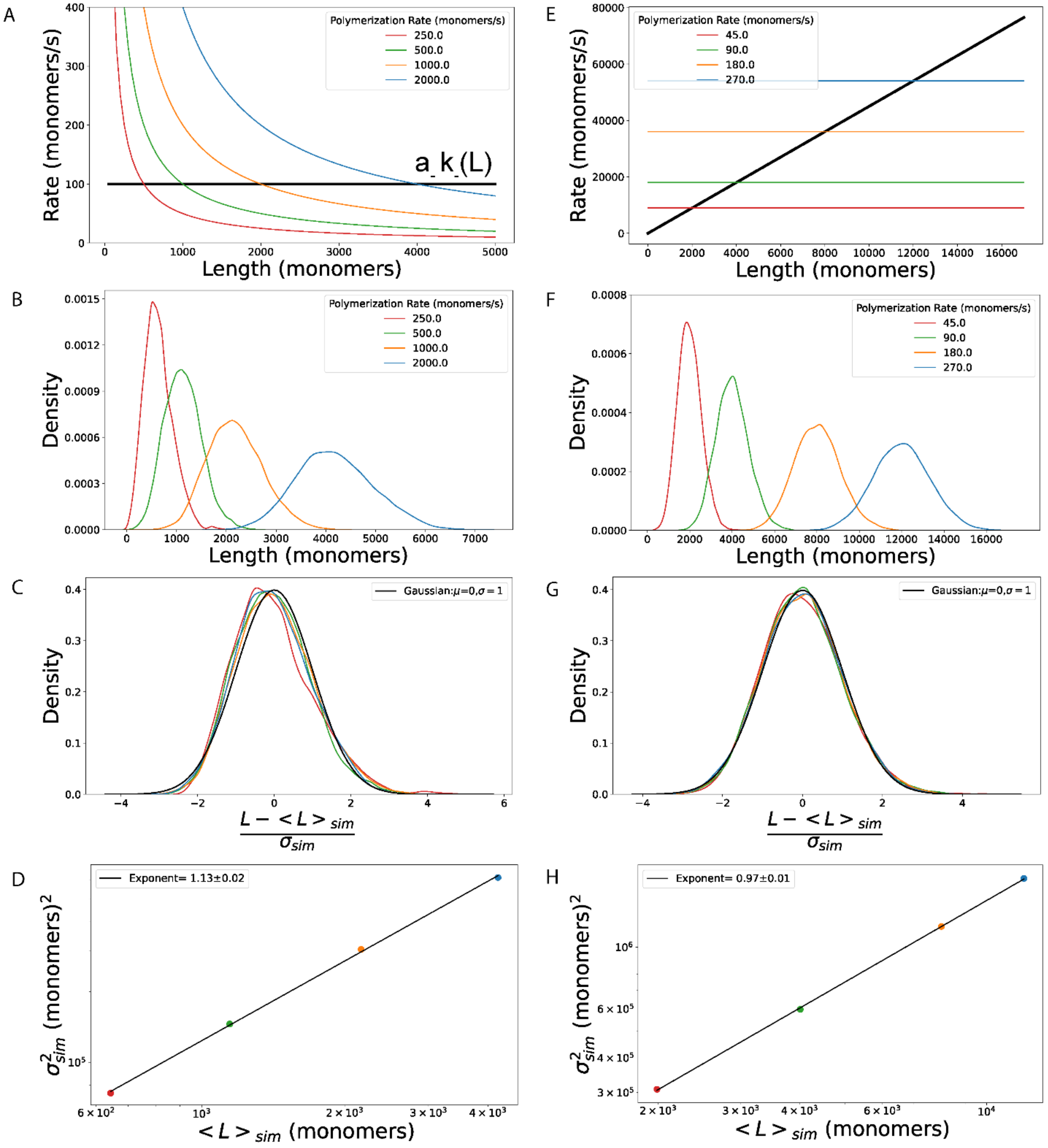


*Supplementary Figure 2:* **Universal length fluctuations in balance point models when $a_+ \neq a_-$:** (A-D) Results for a balance point model with a length dependent polymerization rate, $a_+\kappa_+/L$, and length-independent depolymerization rate $a_-k_-$. Different steady state lengths, $L^*$, are achieved by tuning the polymerization rate parameter, $\kappa_+$: $\kappa_+ = 250\ monomers\ s^{-1}$ (red) ($L^* = 500\ monomers$); $\kappa_+ = 500\ monomers\ s^{-1}$ (green) ($L^* = 1000\ monomers$); $\kappa_+ = 1000\ monomers\ s^{-1}$ (orange) ($L^* = 2000\ monomers$) ; $\kappa_+ = 2000\ monomers\ s^{-1}$ (blue) ($L^* = 4000\ monomers$). In all simulations the length-independent polymerization rate is $k_- = 1\ s^{-1}$. (B) Steady state length distributions from stochastic simulations for different values of $\kappa_+$. (C) The length distributions from (B) collapse to a Gaussian distribution centered around zero with a standard deviation of one, when the lengths are rescaled by the mean and standard deviation of each individual length distribution. (D) The variance of the length distributions scales linearly with the mean length (error bars are standard deviations). (E-H) Results for a balance point model with a length independent polymerization rate, $a_+k_+$, and length-dependent depolymerization rate, $a_-\kappa_-L$. Different steady state lengths, $L^*$ , are achieved by tuning the polymerization rate $k_+$. $k_+ = 45\ s^{-1}$ (red) ($L^* = 2000\ monomers$) ; $k_+ = 90\ s^{-1}$ (green) ($L^* = 4000\ monomers$); $k_+ = 180\ s^{-1}$ (orange) ($L^* = 8000\ monomers$) ; $k_+ = 270\ s^{-1}$ (blue) ($L^* = 12000\ monomers$); In all simulations first order rate of depolymerization, $\kappa_- = 0.045\ monomer^{-1}s^{-1}$. (F) Steady state length distributions for different values of $k_+$, obtained from stochastic simulations (G) The length distributions from (F) collapse to a Gaussian distribution centered around zero with a standard deviation of one, when the lengths are rescaled by the mean and standard deviation of each individual length distribution. (H) The variance of the length distributions scales linearly with the mean length (error bars are standard deviations). In all the simulations the subunit size added in a single polymerization step is $a_+ = 200$ monomers and the subunit size removed in a depolymerization step is $a_- = 100$ monomers.

**Length distribution for severing model of length control:**

We consider a severing model where the filament can undergo polymerization with a length-independent rate, $k_+$ and can be severed with rate $s$ anywhere along the length of the filament with equal probability (Supplementary Figure 3A).

The exact probability distribution of lengths $P(L)$ in steady state, as derived in Mohapatra et al[2], is:

$$P(L) = \frac{L\ s\ k_+^{L-1}}{(k_+ + s)(k_+ + 2s)(k_+ + 3s)\ \dots\ (k_+ + L\ s)} \qquad S29$$

where $L$ is measured in $monomers$, $k_+$ is in units $monomers/second$, and the severing rate parameter $s$ has units $1/second$.

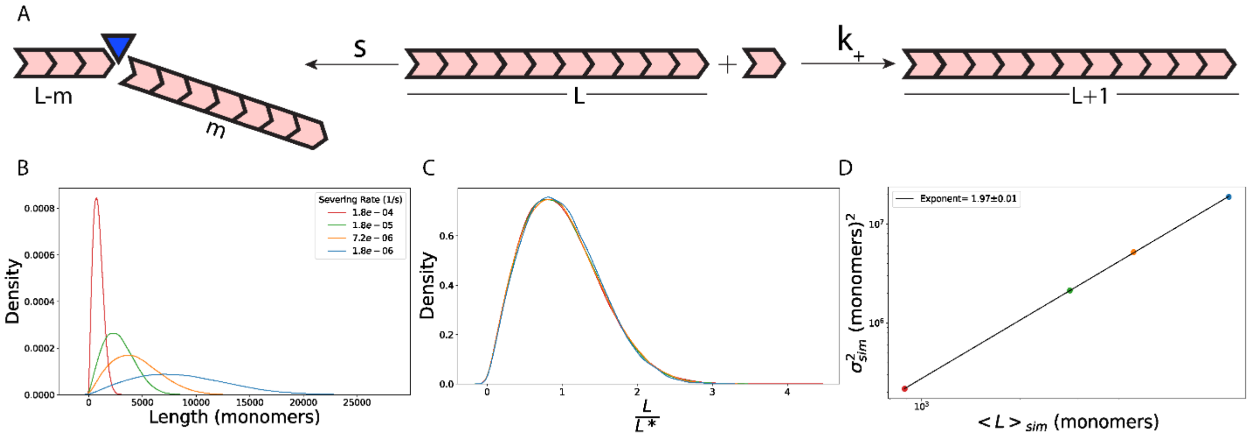


*Supplementary Figure 3:* **Severing model of length control.** *(*A) We consider a model where the filament can grow by addition of a monomer with rate, $k_+$ and is severed with uniform probability along the length of the filament, with a rate $s$. (B) The steady state length can be tuned by changing the severing rate $s$. (Leftmost) The probability distribution of steady state length fluctuations for $s = 1.8 \times 10^{-4}\ monomers/s$ (red) ($L^* = 886\ monomers$); $s = 1.8 \times 10^{-5}\ monomers/s$ (orange) ($L^* = 2800\ monomers$); $s = 7.2 \times 10^{-6}\ monomers/s$ (orange) ($L^* = 4431\ monomers$); and $s = 1.8 \times 10^{-6}\ monomers/s$ (green) ($L^* = 8860\ monomers$). (C) The length distributions from (B) collapse to a universal, Rayleigh distribution centered around one, when the lengths are rescaled by their mean. (D) The variance of the length distributions scales with a power law exponent of $1.97 \pm 0.01$. In all the simulations $k_+ = 90\ monomers\ s^{-1}$ and one monomer is added during a polymerization event.

When the ratio of rate of severing and the rate of polymerization is much less than one, i.e., $^{s}/_{k_+} << 1$ we can approximate the above equation using $\left(1+\ j\frac{s}{k_+}\right)^{-1} \approx e^{-j\,s/k_+}$ :

$$P(L) = \frac{L\,s\,k_+^{L-1}}{k_+^L\left(1+{}^{s}/_{k_+}\right)\left(1+{}^{2s}/_{k_+}\right)\left(1+{}^{3s}/_{k_+}\right)\dots\left(1+{}^{L\,s}/_{k_+}\right)}$$

S30

$$P(L) = \frac{L\,s}{k_+}e^{-\frac{s}{k_+}\sum_{j=1}^{L} j}$$

S31

$$P(L) = \frac{L\,s}{k_+}e^{-\frac{s}{k_+}\frac{L(L+1)}{2}}$$

S32

Assuming $L \gg 1$, namely that the polymer length is much larger than a single subunit, we can further simplify the formula for the distribution to:

$$P(L) = \frac{L\,s}{k_+}\,e^{-\frac{s}{k_+}\frac{L^2}{2}}$$

S33

Using Equation S33, we can compute the mean length:

$$< L >= \frac{\int L\,P(L)dL}{\int P(L)dL} = \sqrt{\frac{\pi}{2\alpha}}$$

S34

where $\alpha \equiv \frac{s}{k_+}$. The variance of the distribution is:

$$Var(L) = \left(\frac{4}{\pi} - 1\right) < L >^2$$

S35

The variance of the distribution grows as the square of the steady state length.

A simple argument based on the balance point result for the variance can be made to get at this result. Namely, since there is equal probability of severing anywhere along the length of the filament the average size of the removed piece ($a$), in one severing step is of the order $a \sim L^*$, where $L^*$ is the length in steady state. Since for the balance point model $Var(L) \sim\ aL^*$ we should expect $Var(L) \sim L^{*2}$ for the severing model.

We performed stochastic simulations of the severing model and found them to be consistent with the analytical results for the steady state length distribution and the scaling of the variance of the distribution with the mean length (Supplementary Figure 3).

**Distribution of bundle lengths when individual filament lengths are exponentially distributed:**

We consider a bundle of parallel filaments whose lengths ($l_i$) are sampled from an exponential distribution:

$$p(l_i) = \frac{1}{< l >} e^{-l_i/_{<l>}} ,$$

*S36*

where $< l >$ is the average length of filaments within a bundle.

The probability that a filament has a length, $l_i < L$ is given by the cumulative distribution function:

$$F(L) = \int_0^L p(t)dt = 1 - e^{-\frac{L}{<l>}}$$

*S37*

The length of the N filaments within the bundle are $l_1, l_2, \dots, l_N$ and the probability that all the N filaments have a length that is less than $L$ is given by:

$$F_N(L) = \left(1 - e^{-\frac{L}{<l>}}\right)^N$$

*S38*

From this cumulative probability distribution, we can compute the probability that the longest of the N filaments in the bundle has a length $L$, (which is also, by definition, the length of the bundle)

$$p_N(L) = \frac{d}{dL}\big(F_N(L)\big) = \frac{N}{< l >}\left(1 - e^{-\frac{L}{<l>}}\right)^{N-1} e^{-\frac{L}{<l>}}$$

*S39*

The mean of this distribution is given by

$$< L > = \int_0^\infty L \ p_N(L) \, dL$$

*S40*

This integral can be computed by making the substitution $e^{-\frac{L}{<l>}} = 1 - u$

$$< L > = - N < l > \int_0^1 \ln(1-u) \ (u)^{N-1} du$$

*S41*

After series expanding the logarithmic term and integrating:

$$< L >= N < l > \sum_{k=1}^{\infty} \frac{1}{k(N+k)}$$

*S42*

Using the identity $\frac{N}{k(N+k)} = \frac{1}{k} - \frac{1}{N+k}$ we can compute the infinite series:

$$\sum_{k=1}^{\infty} \frac{\mathrm{N}}{k(N+k)} = \sum_{k=1}^{N} \frac{1}{k} \equiv H_N$$

*S43*

which gives us a simple formula for the average bundle length

$$< L >= H_N < l > .$$

*S44*

Here, $H_N$ is the Harmonic number, which for large $N$ is given by the approximate formula $H_N \approx \gamma + \ln(N)$, where $\gamma = 0.577$ … is the Euler-Mascheroni constant. Therefore, when the number of filaments in the bundle is large, the average bundle length is

$$< L > \approx < l > (\gamma + \ln(N)).$$

*S45*

To compute the variance of the bundle length we first compute the second moment of the distribution:

$$< L^2 > = \frac{N}{< l >} \int_0^{\infty} L^2 \left(1 - e^{-\frac{L}{<l>}}\right)^{N-1} e^{-\frac{L}{<l>}} \, dL.$$

*S46*

With the help from Wolfram alpha to compute the integral, we find

$$< L^2 > = N < l >^2 \left[\frac{6(H_N)^2 - 6\Psi^{(1)}(N+1) + \pi^2}{6N}\right]$$

*S47*

where $H_N$ is the Harmonic number and $\Psi^{(1)}$ is the first order Polygamma function. For large $N$, this reduces to

$$var(L) = < l >^2 \left[\frac{\pi^2 - 6\Psi^{(1)}(N+1)}{6}\right].$$

*S48*

The function $\Psi^{(1)}(N)$ is much less than one for typical values for the number of filaments in a parallel actin bundle; for $N = 5$ $\left(\Psi^{(1)}(6) \approx 0.18\right)$, while for $N = 1000$ $\left(\Psi^{(1)}(1001) \approx 0.001\right)$. Therefore,

$\Psi^{(1)}(N+1)$ can be ignored when compared to $\pi^2 \approx 9$ in Equation S48, and this leads to the expression:

$$var(L) \approx < l >^2 \frac{\pi^2}{6} .$$

S49

Using Equations S45 and S49 we obtain an expression for the variance of the bundle length in terms of its mean:

$$var(L) = < L >^2 \frac{\pi^2}{6(\gamma + lnN)^2} .$$

S50

**Estimating the number of filaments from the bundle model:**

Equation S51 makes a very sharp prediction, namely when plotting the log of variance of the bundle length as a function of the log of the mean bundle length we expect a straight line with slope 2 and intercept $log_{10} \frac{\pi^2}{6(\gamma+lnN)^2}$ .

The slope we obtain from the fit in Figure 3E is $2.09 \pm 0.1$ (mean ± SEM) which is close to the predicted value of 2; the 95% confidence interval for the fitted slope is $1.9 - 2.3$. The intercept of the fitted line to the data in Figure 3E is $-1.3 \ \pm 0.06$ and the 95% confidence interval is given by $-1.42$ to $-1.18$. From this we can estimate the number of filaments to be between 80 and 400 which compares favorably to the typical number of filaments in the different actin structures that were analyzed. For actin cables the number of filaments $5 - 10$[3], for microvilli it is $10 - 50$[4], for stereocilia it is $100 - 900$[5], and for filopodia it is $10 - 30$[6].

**Distribution of bundle lengths when individual filament lengths are controlled by severing:**

The probability distribution of individual filament lengths ($l_i$) that undergo severing dynamics is given by Equation S33. The average filament length for this distribution is $< l > = \int_0^\infty t\, p(t)dt \ = \sqrt{\frac{\pi}{4}}\, \lambda$ and the variance for the distribution is, $var(l_i) = \lambda^2$, where $\lambda = \sqrt{\frac{2k_+}{s}}$.

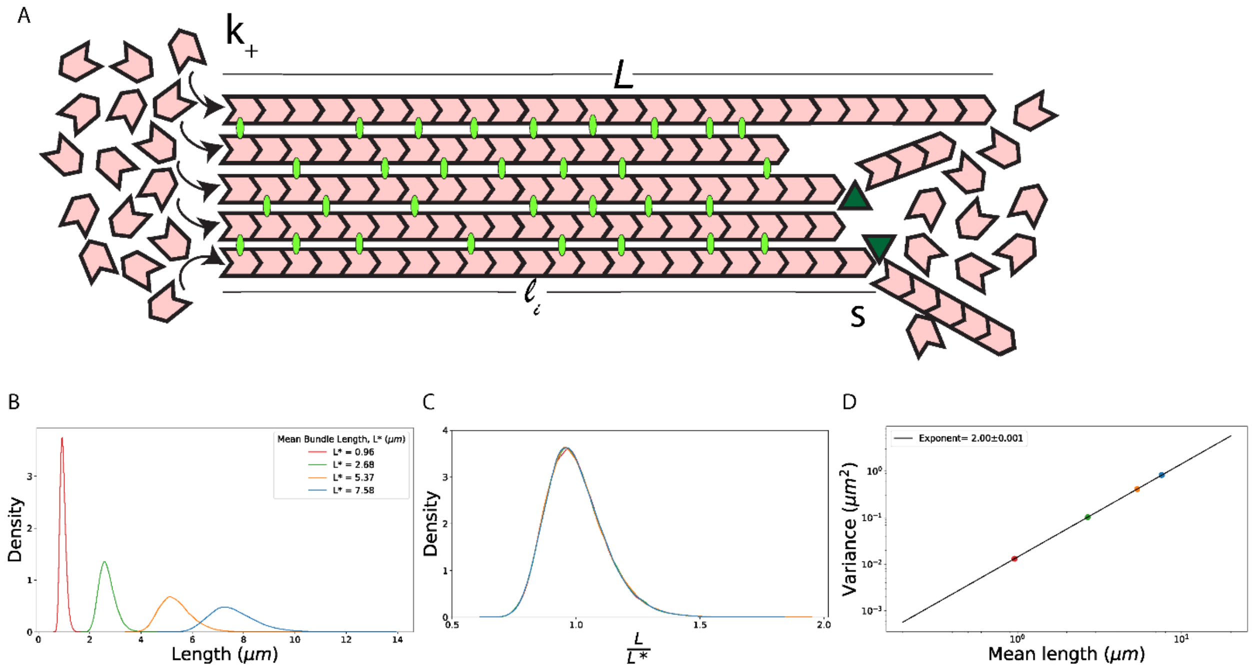


*Supplementary Figure 4:* **Length of a bundle where individual filament lengths controlled by severing**: (A) We consider parallel actin bundles to be made of $N$ actin filaments with lengths $l_1, l_2, \dots, l_N$. Each filament undergoes length changes with a rate of assembly, $k_+$ and can undergo severing with a rate $s$ along the length of the filament. This model gives rise to a Rayleigh distribution of individual filament lengths, $P(l_i) = \frac{l_i s}{k_+} e^{-\frac{s}{k_+}\frac{l_i^2}{2}}$. We define the length of the bundle as the maximum of filament lengths, $L = max\,\{l_1, l_2, \dots, l_N\}$. (B) In simulations random numbers that follow a Rayleigh distribution are generated. These randomly generated numbers are the lengths of the individual filaments within a bundle, $l_i$. With $N = 100$ we find the maximum length of the filaments within a bundle to plot the distribution of bundle lengths. Tuning the severing rate $s$, changes the mean of the distribution of individual filament lengths, $l_i$ given by $< l_i > = \sqrt{(\frac{\pi k_+}{2s})}$, changing the mean of bundle lengths, $L^*$. Here $k_+ \; = \; 0.3\; \mu m/s$ and $< l_i > = \; 0.38\; \mu m$ for $s = \frac{3.33}{\mu m\; s}$ to get $L^* \; = \; 0.96\; \mu m$ (red), $< l_i > = \; 1.05\; \mu m$ for $s = \frac{0.43}{\mu m\; s}$ to get $L^* \; = \; 2.68\; \mu m$ (green), $< l_i > = \; 2.1\; \mu m$ for $s = \frac{1.07}{\mu m\; s}$ to get $L^* \; = \; 5.37\; \mu m$ (orange), $< l_i > = \; 2.97\; \mu m$ for $s = \frac{0.05}{\mu m\; s}$ to get $L^* \; = \; 6.47\; \mu m$ (blue) (C) The distributions of bundle lengths collapse on top of each other when the lengths are normalized by the mean bundle lengths. (D) The log-log plot of the variance as a function of the mean length from the distributions showed in (B) gives the exponent of the power law as $2.00 \pm 0.001$.

To obtain the scaling of the variance of the bundle length distribution with the mean, we make use of mathematical results of the theory or extreme value statistics[7,8]. Namely, the maximum value of $N \gg 1$ random variables, each sampled from a probability distribution whose tail is of the form $p(x) \sim e^{-x^\delta}$, has a Gumbel distribution with scaling parameters $a_N \; = (lnN)^{1/\delta}$ and $b_N \; = \frac{1}{\delta}(lnN)^{-1/\delta - 1}$. In terms of these scaling parameters, the mean of the maximum value is given by $a_N + \gamma b_N$, and the variance is $\frac{\pi^2}{6} b_N^2$, where $\gamma$ is the Euler-Mascheroni constant [9].

Applying this general result to our case, with $x = l/\lambda$ and $\delta = 2$, we arrive at the mean for $L = max_i\{l_i\}$

$$< L > \approx \lambda\sqrt{lnN} \; = \sqrt{\frac{4\; lnN}{\pi}} \; < l >$$

while the variance of the bundle length is given by the formula

$$var(L) = \frac{\pi^2}{24} \times \frac{\lambda^2}{lnN} = \frac{\pi^2}{24} \times \frac{< L >^2}{(lnN)^2}. \tag{S52}$$

Comparison between the analytic results and simulated bundles generated by sampling individual filament lengths from the steady state length distribution of the severing model of length control are shown in Supplementary figure 4.

In Supplementary Figure 5A and 5B we plot the dependence of the mean bundle length and the variance on the number of filaments in the bundle and the average length of a filament in the bundle. The key observation is that while the bundle length is sensitive to the lengths of the filaments in the bundle, it is only weakly (logarithmically) dependent on the number of filaments in the bundle.

In Supplementary Figure 5C we compare the theoretical predictions of this model to the experimental data by comparing the measured and predicted coefficients of variation as a function of mean bundle length. The data are all consistent with the predicted coefficient of variation for bundles of filaments where the number of filaments varies between 10 and 500, as is the case for microvilli, stereocilia, filopodia and actin cables.

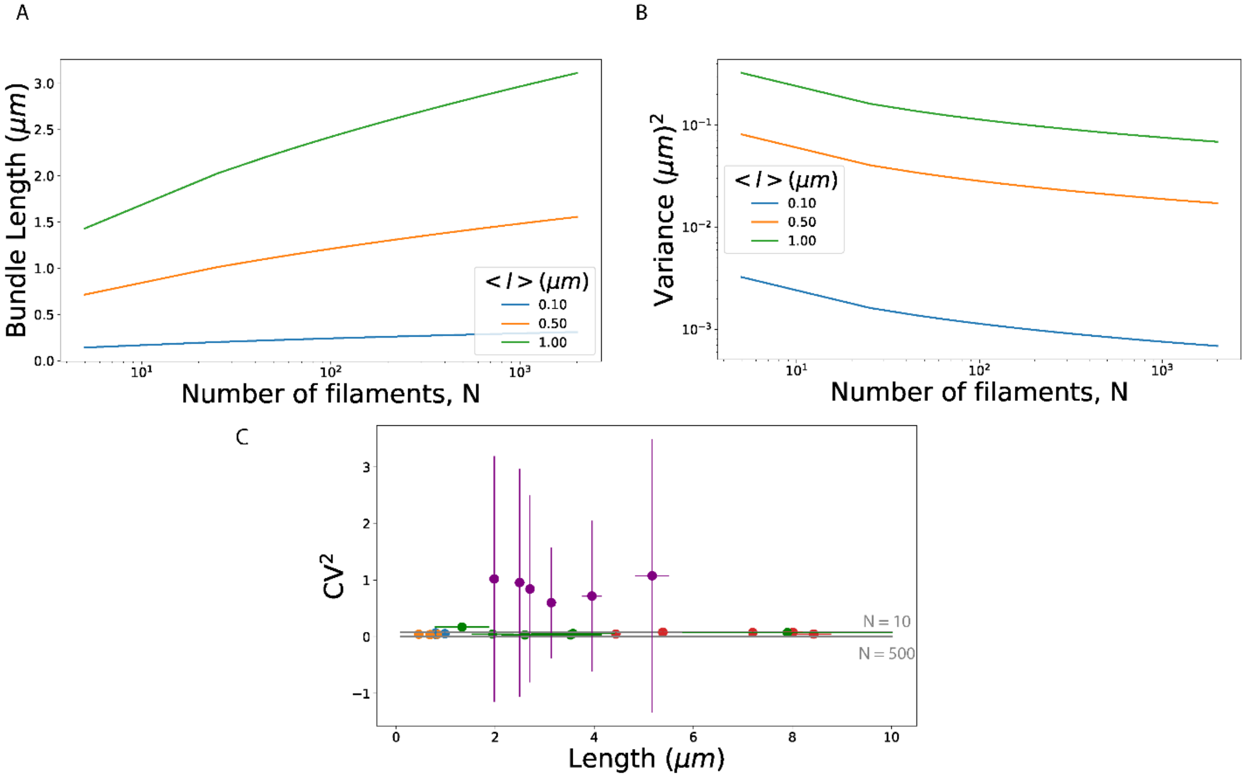


*Supplementary Figure 5:* **Theoretical predictions of the filament-bundle model:** (A) Average length of the bundle, $< L >$ as a function of the number of filaments in the bundle, $N$ for different average lengths of the Rayleigh distributed individual filament within the bundle, $< l >= 0.1\ \mu m$ (blue), $< l >= 0.5\mu m$ (orange), $< l >= 1\mu m$ (green). (B) The variance in bundle lengths as a function of the number of filaments in a bundle, for bundles with filaments sampled from Rayleigh distributions with different means, as in (A). (C) The square of the coefficient of variation is plotted as a function of the mean length for all the data in Figure 3E, following the same color code. The parallel gray lines are predictions from Equation 13 for different number of filaments in the bundle: the top line corresponds to $N = 10$ and the line below corresponds to $N = 500$

### Distribution of bundle lengths when individual filament lengths are controlled by a balance point model:

The probability distribution of individual filaments within a bundle whose lengths are controlled by a balance point model is given by a Gaussian distribution:

$$P(l_i) = \frac{1}{\sqrt{2\pi\sigma^2}} e^{-\frac{1}{2}(\frac{l_i - <l>}{\sigma})^2} \tag{S61}$$

Here the average and variance of the length of filaments within the bundle are $< l >$, and $\sigma^2 \approx < l > a$, respectively; $a$ is the subunit length.

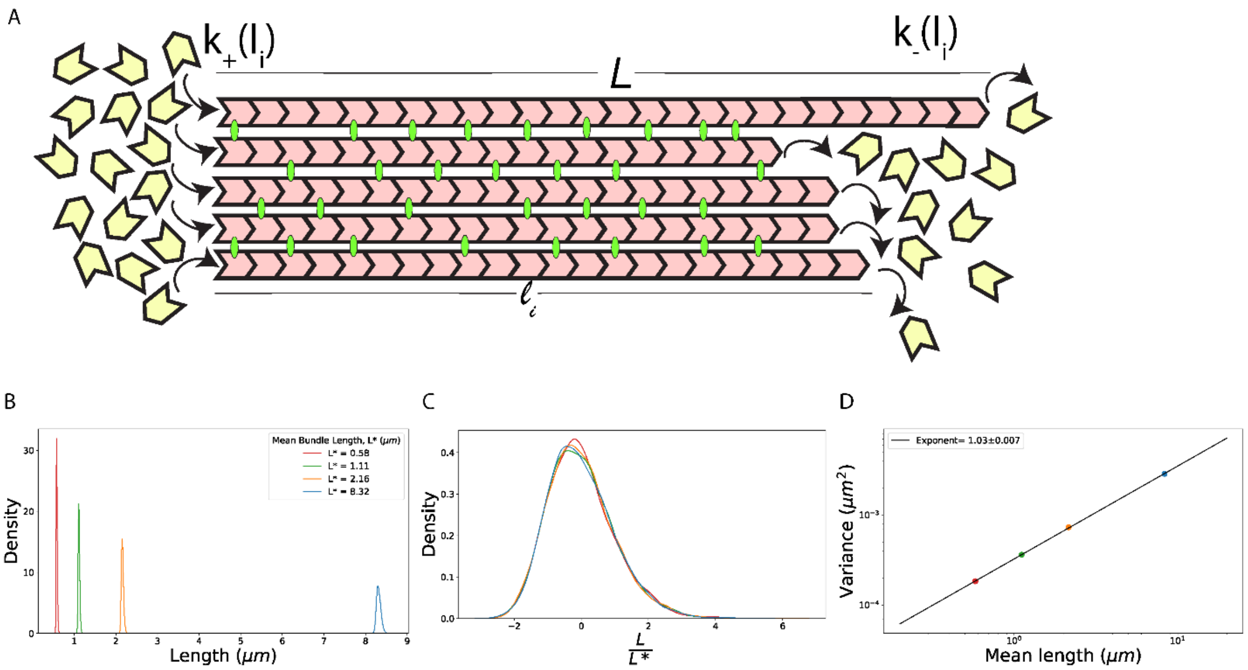


*Supplementary Figure 6:* **Length of a bundle where individual filament length are controlled by the balance point model:** (A) We consider parallel actin bundles to be made of $N$ actin filaments with lengths $l_1, l_2, \ldots, l_N$. Each filament undergoes length changes with a rate of assembly of $k_+(l_i)$ and disassembly of $k_-(l_i)$, where the rates are dependent on the length of the filament. This model leads to a Gaussian distribution of filament lengths, $P(l_i) = \frac{1}{\sqrt{2\pi\sigma}} e^{-\frac{1}{2}(\frac{l_i - <l_i>}{\sigma})^2}$. We define the length of the bundle as the maximum of filament lengths, $L = max\ \{l_1, l_2, \ldots, l_N\}$. (B) In simulations random numbers that are Gaussian distributed are generated. These are the lengths of the individual filaments within a bundle, $l_i$. With $N = 25$ we find the maximum length of the filaments within a bundle to plot the distribution of bundle lengths. By changing the mean, $< l_i >$ and standard deviation, $\sigma$ of the Gaussian distribution we can change the mean bundle length. Here, $< l_i >= 0.5\ \mu m$ and $\sigma = 0.03\ \mu m$ gives $L^* = 0.56\ \mu m$ (red), $< l_i >= 1.0\ \mu m$ and $\sigma = 0.04\ \mu m$ gives $L^* = 1.09\ \mu m$ (green), $< l_i >= 2.0\ \mu m$ and $\sigma = 0.06\ \mu m$ gives $L^* = 2.12\ \mu m$ (orange), $< l_i >= 8.0\ \mu m$ and $\sigma = 0.13\ \mu m$ gives $L^* = 8.25\ \mu m$ (blue) (C) The distributions of bundle length collapse on top of each other when the lengths are normalized by the mean and standard deviation of bundle lengths. (D) The log-log plot of the variance as a function of the steady state length has a slope of 1.

The statistics of the longest filament in the bundle are characterized by the scale parameters $a_N$ and $b_N$ given by[8],

$$a_N = \sqrt{2\ln N} - \frac{\ln(\ln N)}{2\sqrt{2\ln N}} \approx \sqrt{2\ln N}$$

S62

$$b_N = \frac{1}{a_N}$$

S63

The average of the extreme value i.e., the average length of the bundle, $< L >$ is given by[9]:

$$< \frac{L - < l >}{\sigma} > = a_N + \gamma b_N$$

S64

$$< L > = < l > + \sigma \left(\frac{2\ln N + \gamma}{\sqrt{2\ln N}}\right)$$

S65

The variance of the extreme value is given by[9]:

$$var(L) \approx \frac{\pi^2}{12\ln N} \, a < L >$$

S66

The analytic results above are compared to the results from simulated bundles as shown in Supplementary Figure 6. The bundles are made of $N = 25$ filaments whose lengths are samples from a Gaussian distribution with a variance that is proportional to the mean filament length, as is the case for a balance point model.

**Tapering of bundle whose individual filament lengths are sampled from an exponential distribution:**

In the bundled filament model, filaments within the bundle are of varying lengths. This model therefore predicts a thickness profile for the actin structures, $T(x)$. The thickness profile is the number of filaments in the bundle at a given distance $x$ from the growing end, where the filaments within the bundle are polymerized.

The number of filaments within the bundle that have lengths greater than $x$ is given by,

$$T(x) = p(l > x) \times N$$

S67

where $N$ is the number of filaments in the bundle and $p(l)$ is the length distribution of filaments within the bundle which is assumed to be exponential for this derivation.

$$T(x) = N \int_x^\infty \frac{1}{<l>} e^{-l/<l>} \; = \; N\, e^{-x/<l>}$$



So, an exponential length distribution of filaments within the bundle leads to an exponential decrease in the bundle thickness as we move away from the end at which the bundle is polymerized.

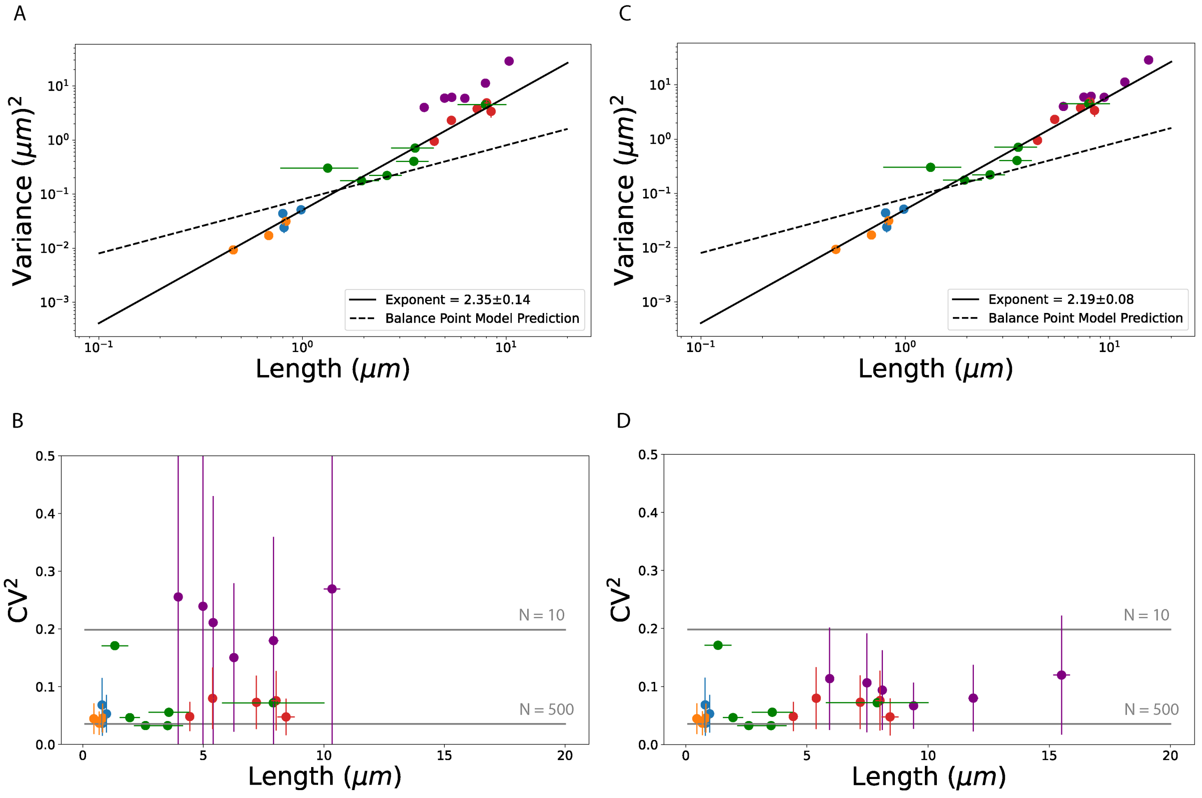


*Supplementary Figure 7:* ***Filopodial actin bundles with lengths modeled as being two or three times longer than the reported lengths of filopodia.*** *(A) Lengths of filopodial actin bundles are assumed to be twice the measured filopodia lengths, which are the ones plotted in Figure 3E. (B) Coefficient of variation for actin filament lengths, with actin bundles in filopodia assumed to be twice the reported filopodia length (C) and (D) same as (A) and (B) but assuming that the lengths of the actin bundles supporting the filopodial membrane protrusions are three times the length of the measured protrusions.*

**Possible explanation for anomalously large variance of filopodia lengths:**

Our analysis of published data for filopodial lengths, while showing the same scaling of variance with the mean as the other filamentous actin structures, show a clear deviation from the universal curve; see Figure 3D-E in the main text. In other words, the coefficient of variation of filopodia lengths is greater than the expected value from the bundled filaments model, which is in quantitative agreement with the variance of the lengths of all the other actin structures we have data for.

One possible explanation for this discrepancy is that the reported filopodia lengths are not of the actin filament bundles that support filopodia but are the lengths of the membrane protrusion they produce. Examination of electron microscopy images of filopodia suggests that the protrusion length is typically considerably smaller than the length of the actin bundle that supports the protrusion. We tested this idea by replotting the variance versus the average length of filopodia assuming that the filopodia lengths are longer than reported by a factor of 2 or 3. We see in Supplementary Figure 7 that a factor of 3 does a good job of accounting for the deviation of the filopodia data from the universal scaling curve. Clearly more experiments must be done to test this hypothesis regarding the deviation of the filopodia length data from the universal curve in Figure 3E.

**Law of total variance and cell to cell variability of rate parameters:**

A key assumption we make when analyzing experimental data and comparing them to the theoretical predictions of the different models of length control is that the measured population variance is a good proxy for the variance of length one would measure in a single cell over time. This is clearly an approximation since even in an isogenic population of cells we should expect differences in the concentration of different cytoskeletal proteins leading to cell-to-cell variation in rate parameters for assembly and disassembly, which in our theory we have assumed to be a constant. In order to assess possible contributions from this source of cell-to-cell variability of rate parameters we turn to the law of total variance. We also recognize that other sources of cell-to-cell variability are possible which underlies the need for single cell experiments of filament length fluctuations.

The variance in a quantity measured across a population of cells, $\sigma_{pop}^2$, has two contributions: (1) the stochastic contribution arising from the variation seen within a cell, $\sigma_{within\ cell}^2$ and (2) the population contribution arising from cell-to-cell variability, $\sigma_{cell-to-cell}^2$. The first contribution would be the measured variance in the population, if each cell was identical in its chemical composition, with identical rate parameters describing the assembly and disassembly of filaments. The second contribution to the total variance would be the one measured if within each cell the chemistry of filament assembly is so precise so that no fluctuations in length arise due to the assembly process. Even though each cell would assemble a very precise filament in this case, slight differences in each cell's chemical composition would result in filament lengths varying from cell to cell. The mathematical statement of the law of total variance is:

$$\sigma_{pop}^2 = \sigma_{within\ cell}^2 + \sigma_{cell-to-cell}^2$$

*S69*

$$\sigma_{pop}^2 = < var(L_i) > + var(< L_i >)$$

*S70*

To asses how the possible presence of the cell to cell variability of rate parameters can affect the variance in length measured in a cell population we assume that filament lengths within each cell are controlled by a balance point model with a length-independent rate of polymerization, $k_+(L) = k_+$ and length-dependent assembly $k_-(L) = \kappa_- L$. The average length in this scenario is given by $< L > = {k_+}/{\kappa_-}$. The variance of length that one would measure in a single cell is given by the balance point model formula derived above (Equation S16) and is ${k_+}/{\kappa_-} \times a$.

Using these results, we can write the total variance of the length as:

$$\sigma_{pop}^2 = < {k_+}/{\kappa_-} \times a > + var\left({k_+}/{\kappa_-}\right)$$

*S71*

Here we assume that due to some difference in chemistry, the rate parameter $k_+$ varies from cell to cell. If we characterize this variation with a population mean $< k_+ >$ and a variance $\sigma_{k_+}^2$,

$$\sigma_{pop}^2 = \; a \; < k_+ >/\kappa_- \; + \frac{\sigma_{k_+}^2}{\kappa_-^2}$$

*S72*

The first term in this expression is nothing but population mean filament length ($< L >_{pop} = < k_+ > /\kappa_-$) times the subunit length $a$, which is the balance point model result we have used thus far. The second term is a contribution to the variance of the length that comes from the cell-to-cell variability in the rate constant $k_+$.

If we assume that the rate parameter fluctuations are due to fluctuations in the amount of a protein that effects polymerization rate, then we can evoke measurements of cell to cell variability in protein abundance. Most such measurements find that the variance scales with the mean protein abundance. In that case we can expect the second term to have the same scaling with mean as the first term in Equation S72, giving an overall scaling of $\sigma_{pop}^2 \sim < L >_{pop}$ . This is inconsistent with the measured length fluctuations across all the different filamentous actin structures for which we have found data (see Figure 3).

If we assume large variations in the rate parameter, such that its variance scales with mean squared, then Equation S72 can be rewritten as

$$\sigma_{pop}^2 = < L >_{pop} \; a + \frac{f^2 < k_+ >^2}{\kappa_-^2},$$

*S73*

$$\sigma_{pop}^2 = < L >_{pop} \; a + \; f^2 < L >_{pop}^2$$

*S74*

where f is the coefficient of variation for the cell-to-cell variability in the rate parameter $k_+$.

In the scenario where there is no noise in the rate parameter ($f = 0$) we revert to the result in the main text that the variance is proportional to the mean length and to the length of the subunits (Equation S16). In the other limit where $f^2 \; >> a/< L >_{pop}$, we predict a variance that scales with the square of the mean length, namely,

$$\sigma_{pop}^2 \approx \; f^2 < L >_{pop}^2$$

*S75*

While this formula is consistent with experiments, we do not think this to be a likely explanation of the observed scaling for two reasons. One, based on numerous experiments, we do not expect protein abundances to have such large variations from cell to cell to produce similarly large rate constant variations. Two, experiments over very different filamentous structures yield coefficient of variation that are universal, independent of structure (Supplementary Figure 7B). In the bundled filament model this is a simple consequence of extreme value distributions, as discussed in the main text. For the scenario discussed in this section to produce an equivalent result we would have to postulate some universal process that applies to all the different cell types to produce the same coefficient of variation ($f$) in one of the rate parameters controlling assembly. This seems unlikely to us.
